# Spatially Restricted Immune Responses Allow for Root Meristematic Activity During Bacterial Colonisation

**DOI:** 10.1101/2020.08.03.233817

**Authors:** Aurélia Emonet, Feng Zhou, Jordan Vacheron, Clara Margot Heiman, Valérie Dénervaud Tendon, Ka-Wai Ma, Paul Schulze-Lefert, Christoph Keel, Niko Geldner

## Abstract

Plants circumscribe microbe-associated molecular pattern (MAMP)-triggered immune responses to weak points of the roots. This spatially restricted immunity was suggested to avoid constitutive responses to rhizosphere microbiota. To demonstrate its relevance, we combined cell-type specific expression of the plant flagellin receptor (FLS2) with fluorescent defence markers and mapped immune competency at cellular resolution. Our analysis distinguishes cell-autonomous and non-cell autonomous responses and reveals lignification to be tissue-independent, contrasting cell-type specific suberisation. Importantly, our analysis divides the non-responsive meristem into a central zone refractory to FLS2 expression, and a cortex that becomes highly sensitised by FLS2 expression, causing meristem collapse upon MAMP exposure. Meristematic epidermal expression generates super-competent lines that detect native bacterial flagellin and bypass the absence of response to commensals, providing a powerful tool for studying root immunity. Our precise manipulations and read-outs demonstrate incompatibility of meristematic activity and defence and the importance of cell-resolved studies of plant immunity.

## Introduction

Plant roots host a vast range of microorganisms in their rhizosphere. Amongst those, some can act as pathogens, negatively impacting plant growth and reproduction. However, the plant’s sophisticated innate immune system keeps the vast majority of pathogens at bay. This MAMP-triggered immunity (MTI) rests on the recognition of highly conserved microbial molecules, recognised by plasma membrane-localised pattern-recognition receptors (PRRs) (Zipfel, 2008). One of the most investigated MAMPs is a 22 amino acid fragment of the bacterial flagellin protein (flg22). It is detected by the FLAGELLIN SENSING 2 (FLS2) receptor (Felix *et al*., 1999; Gómez-Gómez and Boller, 2000; Gómez-Gómez *et al*., 1999; Zipfel *et al*., 2004) and induces a signalling cascade including ROS production, calcium signalling, MAPKs (MITOGEN-ACTIVATED PROTEIN KINASE) phosphorylation and gene transcription, eventually leading to defence responses, such as callose and lignin deposition or phytoalexin production (Lee *et al*., 2019; Li *et al*., 2016).

Yet, plant PRRs equally perceive MAMPs from commensal or beneficial microbes, which are part of the normal plant rhizosphere. Whereas MTI is associated with growth inhibition (Chinchilla *et al*., 2007; Gómez-Gómez and Boller, 2000), a plethora of publications have established a growth promoting action of the soil microbiome (Berendsen *et al*., 2012). It therefore becomes particularly interesting to understand how roots accommodate a rhizosphere community, while avoiding a constant activation of PRRs and the growth-defence trade-off that comes with it. Many researchers have argued that the growth inhibition can be overcome by the ability of commensal microorganisms to supress plant immunity (Yu *et al*., 2019a). In addition, it was recently shown that the root has an inherently dampened MTI until it encounters damage, which locally boosts immune responsiveness (Zhou *et al*., 2020).

Indeed, root immune responses are generally lower than in the shoot, often because of an absence or low abundance of PRRs (Beck *et al*., 2014; Faulkner and Robatzek, 2012). Interestingly, plants restrict their defence to regions considered vulnerable. These coincide with regions where protective endodermal barriers are absent or broken, such as in the elongation zone and at the lateral root emergence sites. It is also where bacteria are found to preferentially accumulate (Beck *et al*., 2014; Bulgarelli *et al*., 2013; De Coninck *et al*., 2015; Faulkner and Robatzek, 2012; Millet *et al*., 2010; Poncini *et al*., 2017; Zhou *et al*., 2020).

Here, we set out to address the relevance of spatially limited responses. Wyrsch *et al*. (2015) ectopically expressed *FLS2* under tissue-specific promoters and their data suggested that all root tissues were competent to mount an immune response provided that *FLS2* is expressed, although the nature of the tissue had a large influence on the strength of the innate immunity responses. Yet, the immune read-outs used in this work were at whole-plant or organ-level resolution and did not allow the authors to ascertain from which cell-type responses were originating, or whether responses were cell-autonomous, regional or systemic. Specifically, MAMP-induced ROS production, as well as cytosolic calcium increases, are known to act in a paracrine, even systemic fashion (Dubiella *et al*., 2013; Gilroy *et al*., 2014, 2016; Marhavý *et al*., 2019). Calcium waves were reported to initiate in the root elongation zone and to spread across tissues after flg22 treatment (Keinath *et al*., 2015; Stanley *et al*., 2018), opening the possibility that MAMP responses are induced in cell layers far away from the site of perception.

To address this issue, we combined new fluorescent markers lines with cell-type-specific FLS2 receptor lines. These marker lines use a triple mVenus fluorochrome coupled to a nuclear localisation signal (*prom::NLS-3xmVenus*). Combining concatemerisation with nuclear concentration generates high sensitivity and allows for a clear cellular assignment, not achievable with cytosolic, ER or PM-localised markers. These lines now enable us to observe damage and defence responses with cellular resolution, adding a crucial layer of complexity to our analyses (Marhavý *et al*., 2019; Poncini *et al*., 2017; Vermeer *et al*., 2014; Zhou *et al.,* 2020). We also added fluorescence-based markers that have been used for assessing cytosolic calcium changes triggered by flg22 at single cell resolution (Thor and Peiter, 2014).

This has allowed us to manipulate and quantitatively map defence responses at cellular resolution in the root. Our approach revealed the presence of regions refractory to FLS2 presence, as well as others which are super-competent. We show that inappropriate *FLS2* expression has drastic impact on root development, affecting growth, cell wall composition and cell viability. To assess the impact of *FLS2* misexpression in response to natural microbiota, we use our super-competent lines in the presence of commensal bacteria, normally not detected by wild-type plants. We demonstrate stimulation of FLS2 directly by native, bacteria-derived flagellin and reveal the importance of spatial restriction of immune responses in order to adequately balance growth and defence.

## Results

### Tissue-specific expression of FLS2

In order to analyse the ability of the different root tissues to respond to flg22, we used lines expressing *FLS2* under cell-type-specific promoters in an *fls2* mutant background (Wyrsch *et al*., 2015). We selected lines expressing *FLS2-GFP* driven by three different tissue-specific promoters: *WEREWOLF* for epidermis (*WER::FLS2*), *CASPARIAN STRIP DOMAIN PROTEIN 1* for endodermis (*CASP1::FLS2*), and *SHORT-ROOT* for inner cell layers (*SHR::FLS2*). As controls, we monitored FLS2-GFP driven by the constitutive promoter *UBIQUITIN 10 (UBQ10::FLS2)* and by the native *FLS2* promoter (*FLS2::FLS2*). As described previously, endogenous *FLS2* expression was observed principally in the differentiated stele (Beck *et al*., 2014) (Fig.1A) but also weakly in all tissues from the elongation to the differentiated zone, as well as in root cap cells (Zhou *et al*., 2020) (Fig.1B). *WER::FLS2,* by contrast, was strongly expressed in the epidermis of the meristematic zone (Fig.1), as predicted (Lee and Schiefelbein, 1999), with some weak signal in the older cortex (elongation zone) (Fig.S1A). In agreement with its established expression (Benfey *et al*., 1993; Helariutta *et al*., 2000), we detected *SHR::FLS2* in the stele close to the meristem (Fig.1), but also faintly in the neighbouring endodermis, suggesting that either FLS2 proteins or mRNAs move through plasmodesmata (Fig.S1D). *CASP1::FLS2* had the predicted exclusive expression in differentiated endodermis (Fig.1, S1B) and *UB10::FLS2* was detected in all tissues throughout the root, from meristem to differentiation zone (Fig.1, S1C).

**Figure 1:**
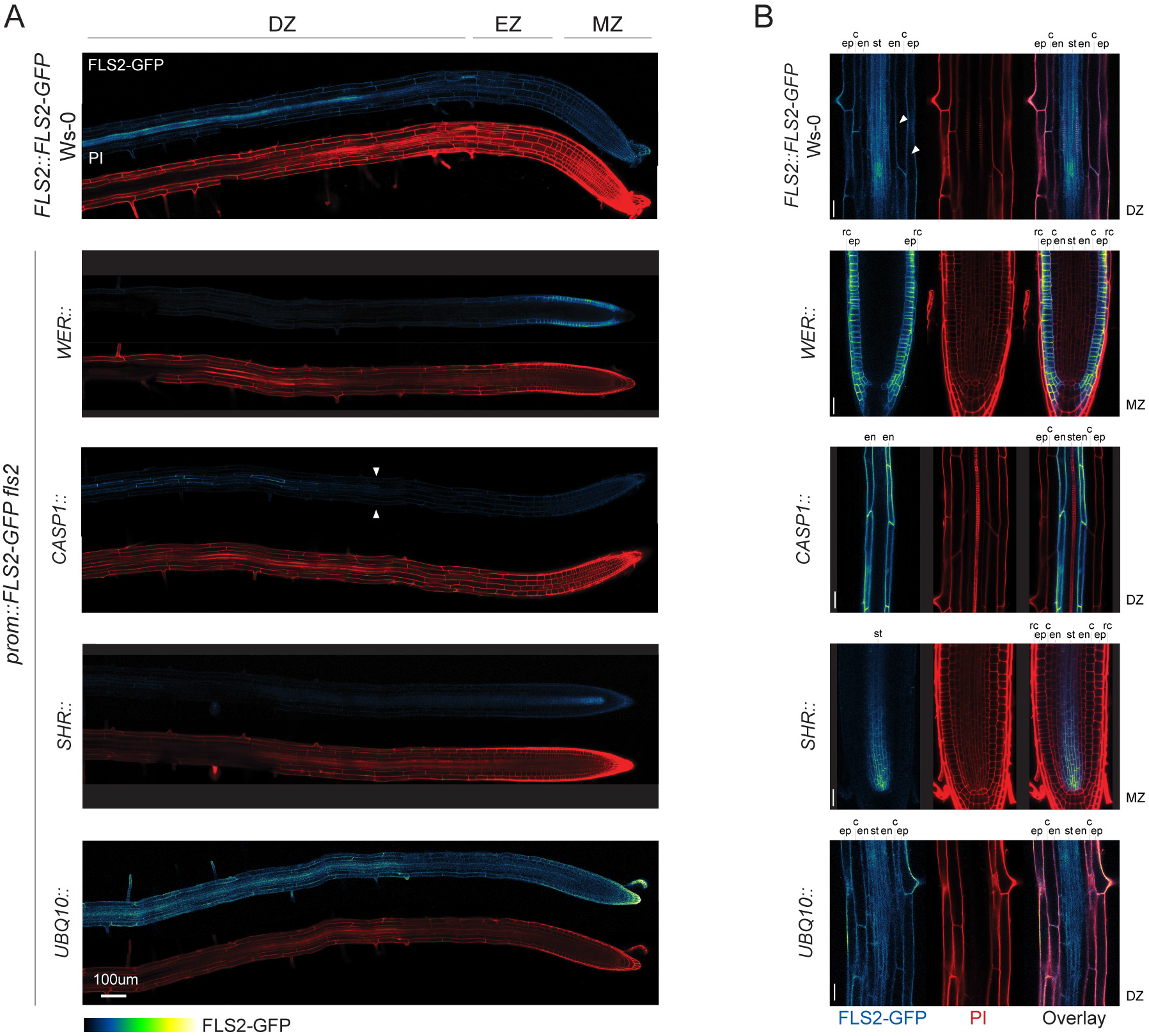
Tissue-specific promoters drive FLS2 receptor expression ectopically. **(A)** Tile scan of *fls2* roots complemented with GFP-tagged FLS2 receptor under epidermal (WER::), endodermal (*CASP1::*), central cylinder (*SHR::*) and ubiquitous (*UBQ10*::) promoters. For comparison, endogenous *FLS2* expression is shown in *FLS2::FLS2-GFP* Ws-0 lines. Root shape is highlighted with PI staining cell wall (PI, red). Scale bar, 100μm. Developmental regions of the roots are labelled: differentiated zone (DZ), elongation zone (EZ), meristematic zone (MZ). **(B)** Close up view of FLS2-GFP expression at selected regions of the complemented lines. *FLS2* driven by its endogenous promoter is expressed in all tissues in the differentiated zone (DZ). Note that in contrast to previous report, low *FLS2* expression is observed in epidermis and cortex (white arrow). In the meristem (MZ), *WER* promoter expresses *FLS2* specifically in epidermis (ep) and root cap (rc), *SHR* promoter in the stele (st) and endodermis (en). In the differentiated zone (DZ), *FLS2* is expressed in all tissues under *UBQ10* promoter, but is restricted to endodermis with *CASP1* promoter. FLS2-GFP (BlueGreen) is co-visualized with PI-stained cell wall (red). Separated and overlaid channels (right column) are presented. Scale bar, 25μm. ep, epidermis; c, cortex; en, endodermis; st, stele; rc, root cap cells.

### Ectopic FLS2 expression alters MTI response patterns

We crossed our selection of FLS2 lines with two typical MTI transcriptional markers, *PEROXIDASE 5 (PER5)* and *MYB DOMAIN PROTEIN 51 (MYB51)*, and generated homozygous lines at all three loci (marker, *prom::FLS2* and *fls2*). As control, we used the two markers in wild-type Col-0 background. Markers were chosen for their strong response to flg22 and their divergent response patterns (Poncini *et al*., 2017; Wyrsch *et al*., 2015; Zhou *et al*., 2020). In addition, we developed a pipeline using tissue-specific quantitative analysis, for measuring and comparing MTI responses in an unbiased fashion (Fig. S2). For this, we additionally introduced ubiquitous nuclear markers (*UBQ10::NLS-mTurquoise* or *UBQ10::NLS-tdTomato*) in all our genotypes, which allows to call all nuclei as separate, individual 3D Regions-of-Interests (ROIs), even those with weak or absent MTI-response. After mock or flg22 treatment and fixation, cell-wall-stained roots were imaged at three different zones of the root: Meristem (MZ), Elongation (EZ) and Differentiation (DZ). Each nucleus was automatically detected as a 3D object and the obtained nuclei object maps were then combined to the cell wall marker channels to manually curate and assign each nucleus to a tissue. Once the selected nuclei were assigned, mean intensity for each cell type per zone per treatment per genotype were calculated and colour coded for the generation of a quantitative MTI-response atlas for each *prom::FLS2* line (Fig.S2, values in Fig.S4).

Our cell-specific quantification and microscopic analysis confirmed that *PER5* is not expressed in absence of flg22 treatment (Fig.2A, 2BC), but that *MYB51* presents a basal, flg22-independent expression in the epidermis and root cap cells of the undifferentiated tissues (MZ and EZ) and in the stele and the cortex of the DZ (Fig.S3A, S3C). In wild-type plants, both MAMP markers are strongly induced in the EZ, recapitulating previous observations (Fig.2A and S3A) (Millet *et al*., 2010; Poncini *et al*., 2017; Zhou *et al*., 2020). Specifically, *PER5* is triggered almost exclusively in the elongating epidermis and root cap cells (Fig.2B, 2C, 2D). *MYB51* induction is restricted to these same tissues close to the meristem, but induction expands to cortex and pericycle cells in the later root (Fig.S3C, S3D).

**Figure 2:**
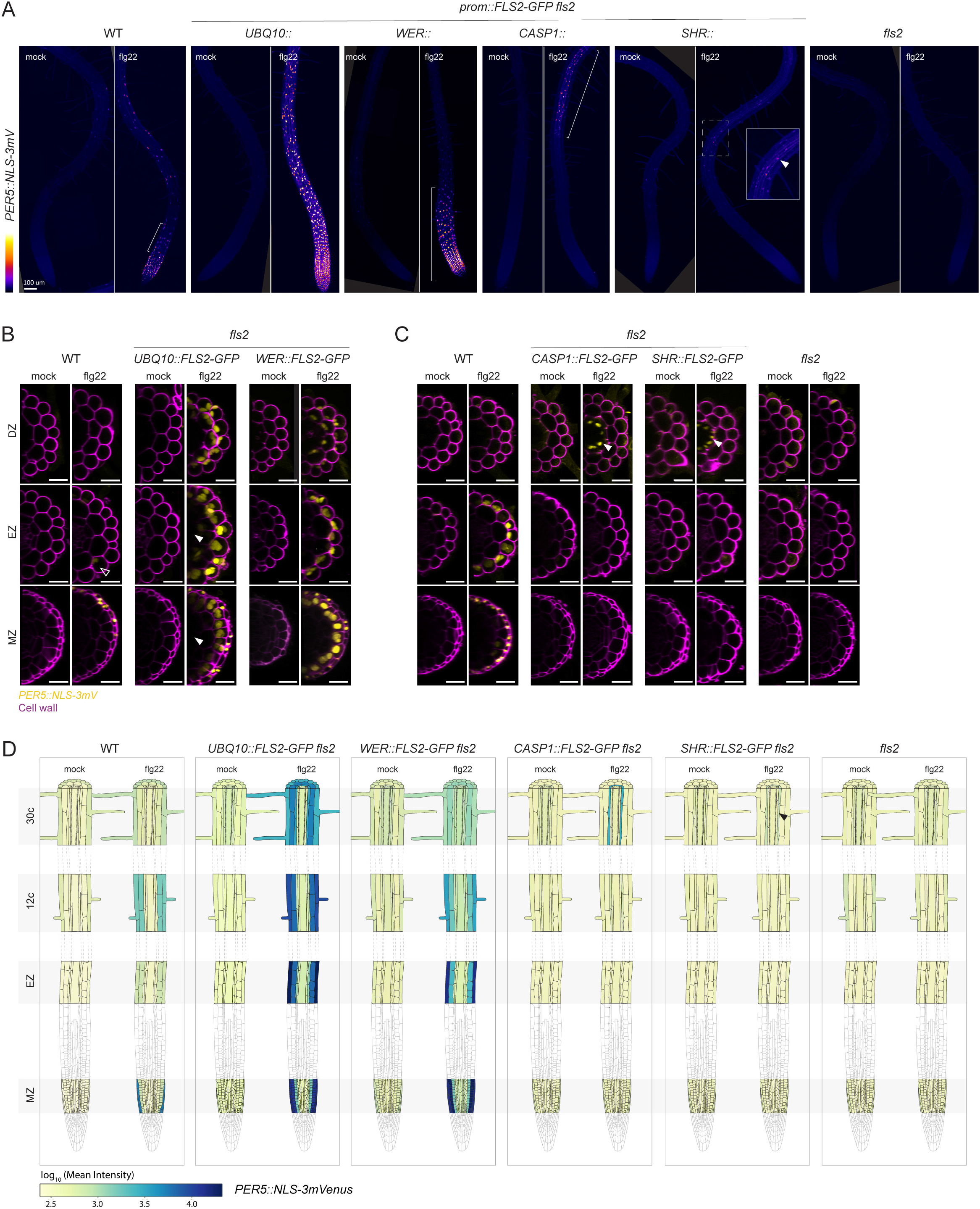
*PER5* marker gene is induced cell-autonomously by flg22 treatment. **(A)** Overview of *PER5::NLS-3mVenus* marker response to flg22 in different *FLS2* recombinant lines. Tile scan images of 1μM flg22 treated plants versus mock. Pictures were taken with similar settings. Settings were always identical between mock and corresponding flg22 treatment. Region of responsiveness is modified by the different expression patterns of *FLS2*. Brackets indicate responsive regions. For *SHR*, close-up view was generated with increased brightness to highlight stellar signal (white arrow). Scale bar, 100μm. **(B)** Maximal projection of transverse sections views of *PER5* expression pattern in UBQ10:: and *WER::FLS2-GFP fls2* compared to WT shown for meristematic zone (MZ), elongation zone (EZ) and differentiated zone (DZ, 30 cells after start of elongation). Seedlings were treated for 24h with 1μM flg22. Note the refractory region in the central cylinder in *UBQ10::FLS2* (white arrows). Nuclear localized mVenus signal (yellow) was co-displayed with propidium iodide cell wall marker (PI, purple). Images were taken with similar settings, but corresponding mock and flg22 treatment pictures for each zone separately always have identical parameters. Note that epidermal signal in flg22-treated wild-type seedlings is faint (EZ, black arrow), due to settings chosen to avoid saturation of signal in the transgenic lines. Compare to Fig.2C, WT. Scale bar, 25μm. **(C)** Maximal projection of transverse section views of *PER5*::*NLS-3mVenus* expression pattern in *CASP1::* and *SHR::FLS2-GFP fls2* as well as WT and *fls2* control. White arrows point at ectopic response in the endodermis. Images were acquired as in Fig.2B., with similar settings between genotypes, but with identical parameter for corresponding mock and flg22 treatment. Pictures were acquired with increased gain compared to Fig.2B due to lower average signal intensity. Scale bar, 25μm. **(D)** Quantitative map of *PER5*::*NLS-3mVenus* responses inferred from tissue specific quantification after 24h treatment with 1μM flg22. Nuclear signals were quantified in ROI delimited with *UBQ10::NLS-mTurquoises2* for all tissue-specific promoter lines, while wild-type (WT) signal was quantified with *UBQ10::NLS-tdTomato* marker. Mean intensity is therefore comparable between *prom::FLS2-GFP fls2* lines, but not to wild-type.

For both markers, changing expression of *FLS2* had an obvious impact on the pattern of responses. Rather than remaining restricted to the elongation zone, *PER5* and *MYB51* induction largely follows the ectopic *FLS2* expression pattern. The defence markers extend to the whole root in *UBQ10::FLS2*, while they are restricted to the DZ or the MZ in *CASP1::FLS2* and *WER::FLS2*, respectively (Fig.2A and S3A). As expected, the *fls2* mutant does not respond to flg22 in any tissue.

*PER5* responds only in the differentiated endodermis in the *CASP1::FLS2* recombinant line, which matches with the very specific expression pattern of *CASP1* promoter. For *WER::FLS2* line, the *PER5* response also follows *FLS2* expression. We could quantify a strong response in root cap cells and the meristematic epidermis, extending until the early DZ, as well as in cortex cells, where we could also detect FLS2 protein (Fig.2B, 2C, 2D, S1A). In contrast to *PER5*, we detected *MYB51* response to flg22 not only in cells expressing *FLS2,* but also some degree of induction in neighbouring cells (Fig. S3B, S3C, S3D). Intensity ratio between flg22 and mock treated plants were calculated and represented graphically in Fig.S5. Non-cell-autonomous responses were obvious for *MYB51* in the DZ of *CASP1::FLS2*. Although *FLS2* is specifically expressed in the endodermis, we could barely detect any *MYB51* responses in this tissue, while the neighbouring stele and cortex cells strongly up-regulated *MYB51* (Fig.S3C, S3D, S5). Similarly, flg22 treatment led to *MYB51* expression not only in the epidermis and cortex, but also in central tissues in *WER::FLS2*. Thus, we concluded that *MYB51* induction by MAMPs is controlled by non-cell autonomous mechanisms, in contrast with the strict cell-autonomy of *PER5* and *FRK1* (this work and Zhou *et al*., 2020).

### FLS2 expression is insufficient to cause flg22-responses in the vascular meristem

Intriguingly, some tissues were also completely refractory to flg22-triggered responses. Despite a clear presence of FLS2 in the vascular meristem (Fig.1B), flg22 treatment did not trigger *PER5* or *MYB51* in *SHR* and *UBQ10::FLS2* lines (Fig.2, S3), except for some weak *MYB51* induction in meristematic pericycle cells in *UBQ10::FLS2* (Fig.S3D, S5). We conclude that flg22-induction of *MYB51* in the pericycle is due to a non-cell autonomous signal from outer cell layers. Thus, central meristematic tissues differ from outer tissue layers in their competence to respond to flg22 in the presence of receptor.

### Ca^2+^ waves are non-cell autonomous responses

Cytosolic Ca^2+^ increases are among the earliest responses upon MAMP perception, preceding transcriptional changes (Jeworutzki *et al*., 2010; Seybold *et al*., 2014). In roots, Ca^2+^ influx after flg22 perception was shown to spread across tissues (Keinath *et al*., 2015). However, since many cells express some degree of *FLS2* in wild-type, it is impossible to dissect to what extent such waves represent a non-cell autonomous propagation of the Ca^2+^ signalling, or are due to flg22 diffusion and direct stimulation of the different tissue layers and regions. We therefore introduced the intensity-based Ca^2+^ reporter *R-GECO1* in our transgenic lines (Keinath *et al*., 2015). We observed in *WER::FLS2*that, like in WT (Movie 1 and 6), calcium signals initiate in the epidermis and spread to inner tissues (Fig.3AB, Movies 2 and 7). Since the receptor was not expressed in central tissues, this clearly demonstrates the non-cell autonomous nature of FLS2-stimulated calcium signalling. This spreading of Ca^2+^ could be observed in all recombinant lines tested, with the intriguing feature that wave direction could be manipulated – *i.e.* in both *CASP1::FLS2* and *SHR::FLS2* lines, the wave started first in the endodermis then spread to outer and inner tissues (Fig.3CD, Movie 8 and 9). Moreover, in these two lines, the wave starts in the differentiated zone rather than in the elongation zone (Movies 3 and 4). When *FLS2* was expressed in all tissues under *UBQ10* promoter, all tissues respond almost simultaneously (Fig.3E, Movie 5 and 10). Taken together, while transcriptional read-outs are largely cell-autonomous, with some degree of tissue-specificity, cytosolic calcium increases represent a non-cell autonomous signalling branch. This implies that even cells that are neither exposed to MAMPs, nor possessing perception capacity, are nonetheless rapidly receiving some sort of stress signal in the form of a calcium wave.

**Figure 3:**
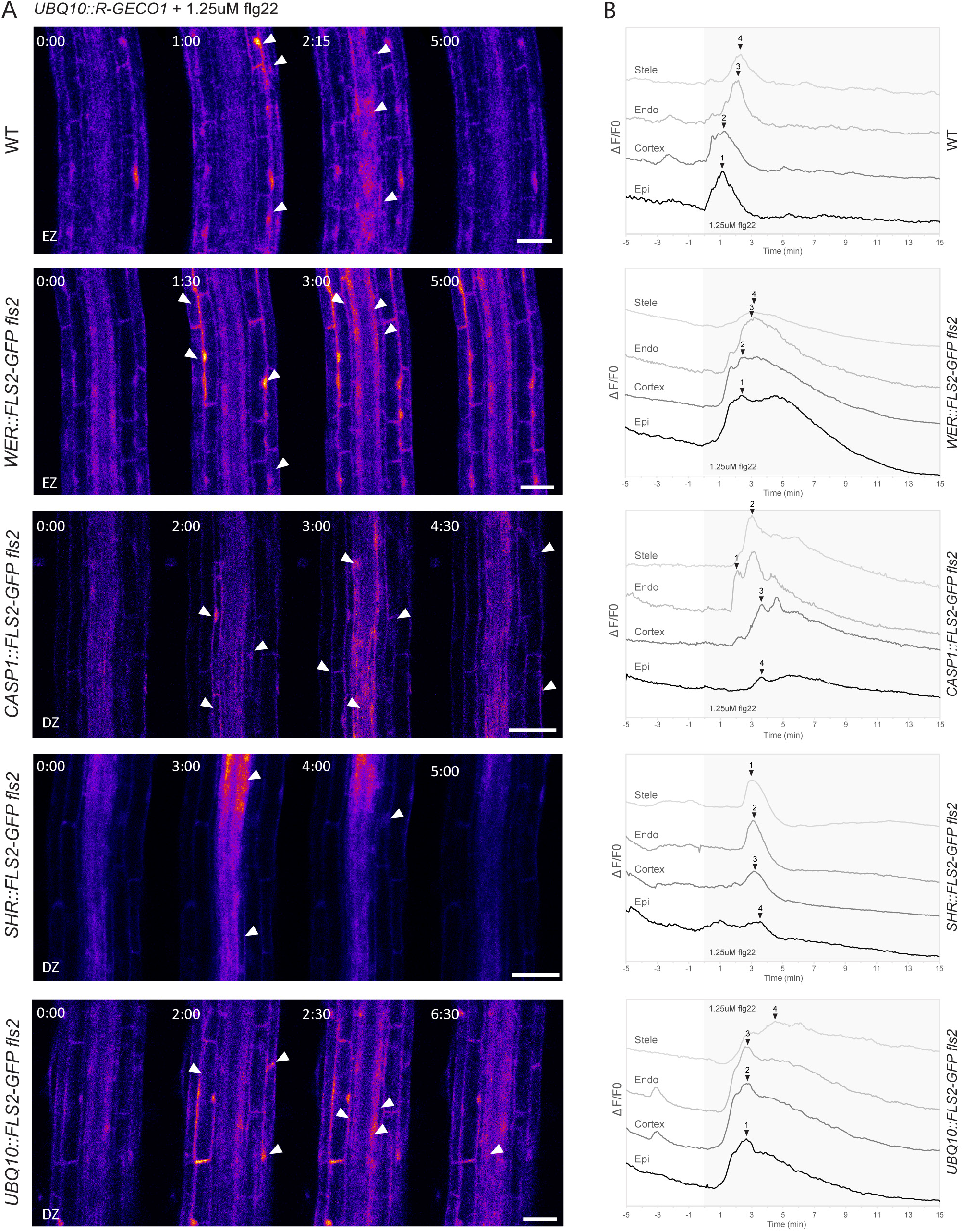
Ca^2+^ waves are non-cell autonomous responses. **(A)** Ca^2+^-dependent signal in the *prom::FLS2-GFP fls2* lines in response to 1.25μM flg22. Time series of *UBQ10::R-GECO1* fluorescence: pictures are longitudinal middle sections of roots at the elongation zone (EZ) or differentiated zone (DZ). Time 0:00 corresponds to the start of flg22 treatment. White arrows point at tissues showing a strong increase in Ca^2+^ content. Scale bar, 25μm. **(B)** Normalized R-GECO1 fluorescence intensity (ΔF/F) measured in tissue-specific ROIs. Values present the dynamics of Ca^2+^ cytosolic concentration in response to flg22 in the root shown in (A) for each tissue type. Black arrows point at the maximum intensity of the trace. Grey background corresponds to flg22 treatment.

### Epidermal meristematic expression of FLS2 leads to flg22 hypersensitivity and meristem collapse

As demonstrated above, *FLS2* ectopic expression can profoundly alter the pattern of immune responses in the root. To test whether this change affects root development, we assessed root length of seedlings transferred on flg22-containing medium. As expected, treated wild-type plants showed only a mild reduction in root length. By contrast, the root length of the constitutive, overexpressing *UBQ10::FLS2* line was strongly reduced with additionally stunted shoot development (Fig.4A and 4B). More surprisingly, a strong root length inhibition was also observed in the *WER::FLS2* line, although this lines expresses *FLS2* only in young epidermal and root cap cells. *SHR::FLS2* and *CASP1::FLS2*, by contrast, showed root growth similar to wild-type.

**Figure 4:**
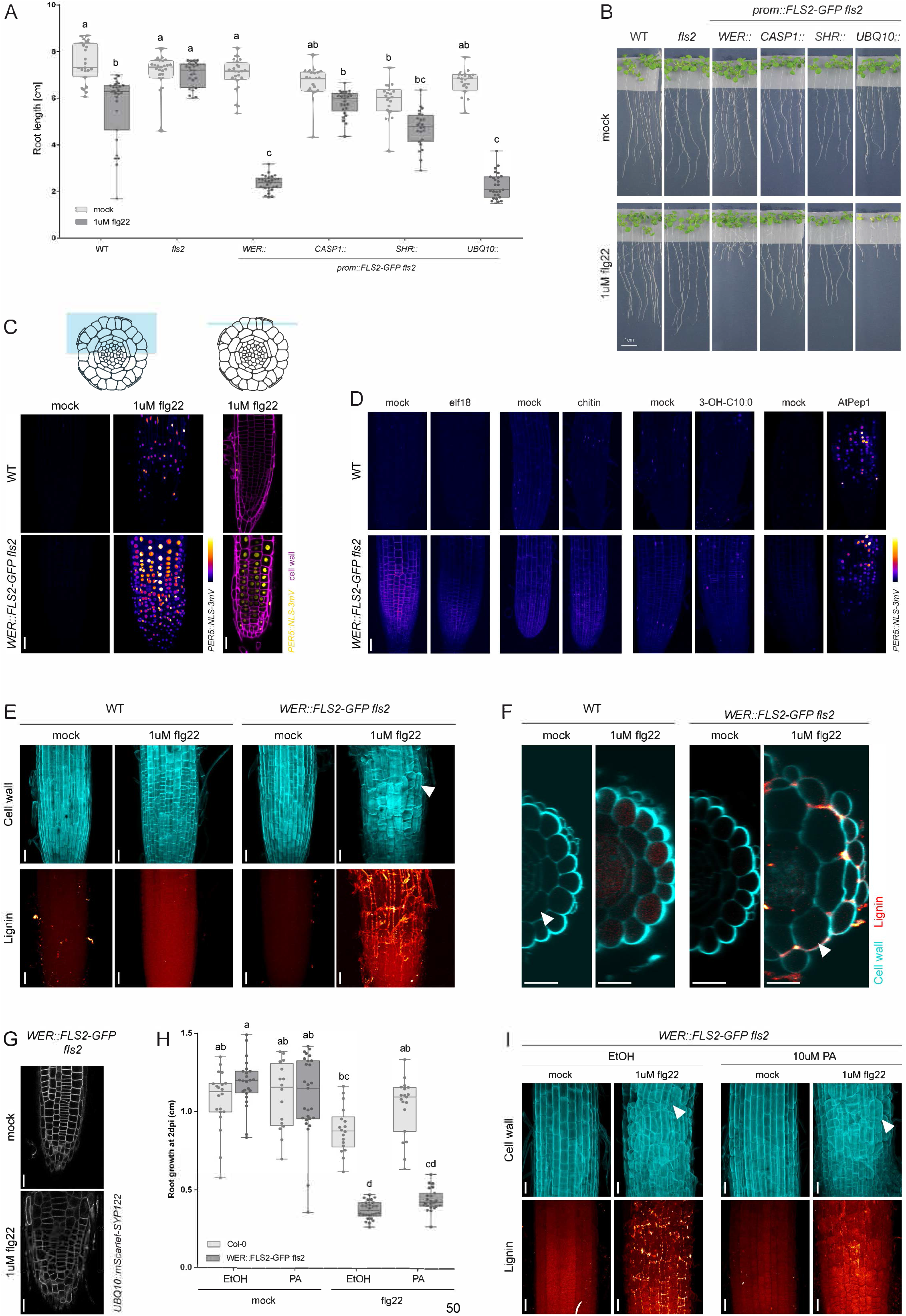
Epidermal meristematic expression of *FLS2* leads to flg22 hypersensitivity and meristem collapse. **(A)** Flg22 treatment increases root growth inhibition in *WER::FLS2* hypersensitive line. Root length quantification of *prom::FLS2-GFP fls2* lines transferred on 1μM flg22 for 6dpi. Boxplot centre represents the median (n=23 to 28 roots). Different letters indicate statistically significant difference between means by Kruskal-Wallis test and Dunn’s multiple comparison. **(B)** Representative pictures of seedlings transferred for 6 days on 1μM flg22. Scale bar, 1cm. **(C)** Flg22 induces strongly *PER5*::*NLS-3mVenus* in the epidermis of *WER::FLS2-GFP fls2* hypersensitive line. On the right, maximum projection of z-stacks taken in root tips of plants treated for 24h with 1μM flg22 or mock. Schematic represents the depth of the z-stack. Pictures were taken with identical settings. Scale bar, 25μm. **(D)** *WER::FLS2-GFP fls2* hypersensitivity is specific to flg22. *WER::FLS2-GFP fls2* and wild-type plants were treated for 24h with either 1μM elf18, 2μg/ml chitin, 1μM 3-OH-C10:0 or 1μM AtPep1. Maximum projection of z-stacks taken in root tips. *PER5* induction is highlighted with mVenus (Fire LUT). Parameters were identical for mock and treatment. Scale bar, 25μm. **(E)** Treatment of *WER::FLS2-GFP fls2* for 2 days with 1μM flg22 induces meristem swelling and lignin deposition. Upper panel shows median projection of calcofluor white stained cell wall in the transition zone of the root tip (blue). Note bulky cells of the epidermis (white arrowhead). Lower panel presents maximum projection of lignin deposition stained with basic fuchsin (red). Lignin accumulates between cells only in *WER::FLS2-GFP fls2* after flg22 treatment. Scale bar, 25μm. **(F)** Cross-section of pictures in (E). Cell wall stained with calcofluor white (blue) is co-visualized with lignin stained with basic fuchsin (red). Flg22 treatment induces massive swelling of cortex cells (white arrowheads) only in *WER::FLS2-GFP fls2*. Lignin is principally deposited between epidermal and cortex cells. Epidermal cells are pushed apart by the swelling cortex and are sometimes missing. Scale bar, 25μm. **(G)** Epidermal view of plasma membrane visualized by the construct *UBQ10::mScarlet-SYP122* in *WER::FLS2-GFP fls2*. Cell division is disorganized after 1μM flg22 treatment. Scale bar, 25μm. **(H)** Inhibition of monolignol synthesis does not rescue meristem flg22-driven increased root growth inhibition of *WER::FLS2-GFP fls2*. Root growth measured after overnight pre-treatment with 10uM PA inhibitor followed by 36h 1μM flg22 combined to 10μM PA treatment. Boxplot centre represents the median (16 <= n <= 27). Different letters indicate statistically significant difference (p<0.05) between means by Kruskal-Wallis test and Dunn’s multiple comparison. **(I)** Flg22 induces meristem swelling despite inhibition of monolignol by PA treatment. Pictures taken from samples quantified in (H). Upper panel shows median projection of calcofluor white stained cell wall in the transition zone of root tip (blue). Lower panel presents maximum projection of lignin deposition stained with basic fuchsin (red). White arrowheads points at examples of bulky cells. Scale bar, 25μm.

In order to precisely identify the tissue responsible for root growth inhibition, we generated two additional *prom::FLS2* lines using the *RCH1 (RECOGNITION OF C.HIGGINSIANUM)* and *PRP3 (PROLINE-RICH PROTEIN 3)* promoters (Marquès-Bueno *et al*., 2015). *RCH1* is expressed in the whole meristem, while *PRP3* is expressed strongly in differentiating root hair cells (Fig.S1E). While *PER5* induction followed the expression of *FLS2* in both lines (Fig. S1GH), only *RCH1::FLS2* presents an increased root growth inhibition (Fig.S1F), whereas *PRP3::FLS2* responds as wild-type (Fig.S1I). Therefore, we conclude that it is the expression of *FLS2* in meristematic epidermal cell layers that causes hypersensitive root growth inhibition in response to flg22. Indeed, when comparing the pattern of *PER5* expression between wild-type and *WER::FLS2* at high resolution, it is evident that only the meristematic epidermal cells show strong *PER5* induction in *WER::FLS2*, whereas root cap cells show flg22 responsiveness in both lines. This suggests that MTI in epidermal cells are the cause of super-competent response (Fig. 4C). Importantly, neither the MAMPs elf18, chitin or the LPS fragment 3-OH-C10:0, nor Atpep1 showed enhanced *PER5* in *WER::FLS2* (Fig.4D). This demonstrates that ectopic *FLS2* expression does not cause a global upregulation of responsiveness to MAMPs, but specifically affect flg22 signalling.

Interestingly, treatment of *WER::FLS2* super-competent line with flg22 induces profound morphological changes in the root, not observed in wild-type. After two days of treatment, cells reaching the transition zone start to swell and division patterns become disorganized, giving rise to bulky meristem shapes (Fig.4E, upper panel, 4G). Virtual cross-sections revealed that cortex cells expand tremendously, dislocating epidermal cells (Fig.4F). Thus, precise spatial regulation of *FLS2* expression levels is necessary to avoid severe growth inhibition caused by flg22-induced disorganized cell expansion in the meristem.

### FLS2 ectopic expression leads to cell-autonomous, flg22-triggered lignin deposition

MTI is known to modify cell wall composition, such as callose deposition or lignification (Chezem *et al*., 2017; Lee *et al*., 2019; Millet *et al*., 2010). Indeed, lignin or suberin deposition are long-known damage and immune responses (Bernards, 2002; Hijwegen, 1963; Kamula *et al*., 1994; Messner and Boll, 1993; Ranathunge *et al*., 2008; Thomas *et al*., 2007), but have not been widely adopted in modern studies on MTI (Lange *et al*., 1995; Mandal and Mitra, 2007), see (Chezem *et al*., 2017; Lee *et al*., 2019) for exceptions.

Interestingly, we found that flg22 treatment induced strong lignification from transition to differentiated zone in WER::FLS2 (Fig. 4E and S6A). Lignin was deposited between epidermis and cortex cells, mainly at the corners (Fig. 4F). In younger regions, lignin was also found between epidermis and root cap cells. All other recombinant lines also showed lignin deposition following their respective *FLS2* expression pattern, except in the stele, matching the absence of *PER5* response in these tissues (Fig.S6B). Interestingly, no lignin deposition could be observed in flg22-treated wild-type roots (Fig.4E, S6), fitting with previous reports (Chezem *et al*. (2017). It is intriguing to speculate that *PER5*, ROS-production and other flg22-responsive genes, categorised as “oxidative stress” response genes (Tognolli *et al*., 2002), are actually part of a lignification response that stays below a productive threshold in wild-type, but pivots into a full lignification upon flg22-stimulation of FLS2 overexpression lines.

The stronger root growth inhibition observed in the super-competent *WER::FLS2* line could be due to the impact of lignin deposition in the transition zone. To test if cell wall reinforcement by lignin prevents cell division and elongation, we inhibited lignin formation with the monolignol synthesis inhibitor piperonylic acid (PA), expecting to restore root growth (Fig. 4I). Nevertheless, even in the absence of lignin, *WER::FLS2* still showed root meristem collapse and stronger RGI than wild-type (Fig. 4H).

### Suberin lamellae deposition after flg22 treatment is an endodermis-specific response

Ectopic lignin deposition occurs in the endodermis as a compensatory mechanism for deficient Casparian strip formation, and is often followed by suberin lamellae deposition (Doblas *et al*., 2017). We wanted to assess whether the overactivation of MTI could trigger the deposition of suberin in cells expressing *FLS2*. In wild-type untreated plants, suberin is usually present in the endodermis only, starting in the late differentiation zone by patches (“patchy zone”), then progressing to a fully suberized zone (Andersen *et al*., 2015, 2018).

In wild-type, suberin was not induced by flg22 (Fig.5AB). In contrast, lines expressing *FLS2* in the endodermis, such as *CASP1::, SHR::* and *UBQ10::FLS2*, showed increased endodermal suberization, leading to a complete disappearance of the patchy zone (Fig. 5B). Earlier suberisation is not simply due to earlier differentiation of endodermal cells due to growth arrest, since *WER::FLS2* still conserved a normal proportion of patchy and suberized zone despite its shorter root length. Reversely, *CASP1::* and *SHR::FLS2* root growth was not affected by flg22, but suberin formed nevertheless much earlier. Therefore, flg22 can induce suberization only when expressed in the endodermis. This endodermis-specific suberisation is a nice demonstration of a flg22 response that only occurs in a specific cellular context.

**Figure 5:**
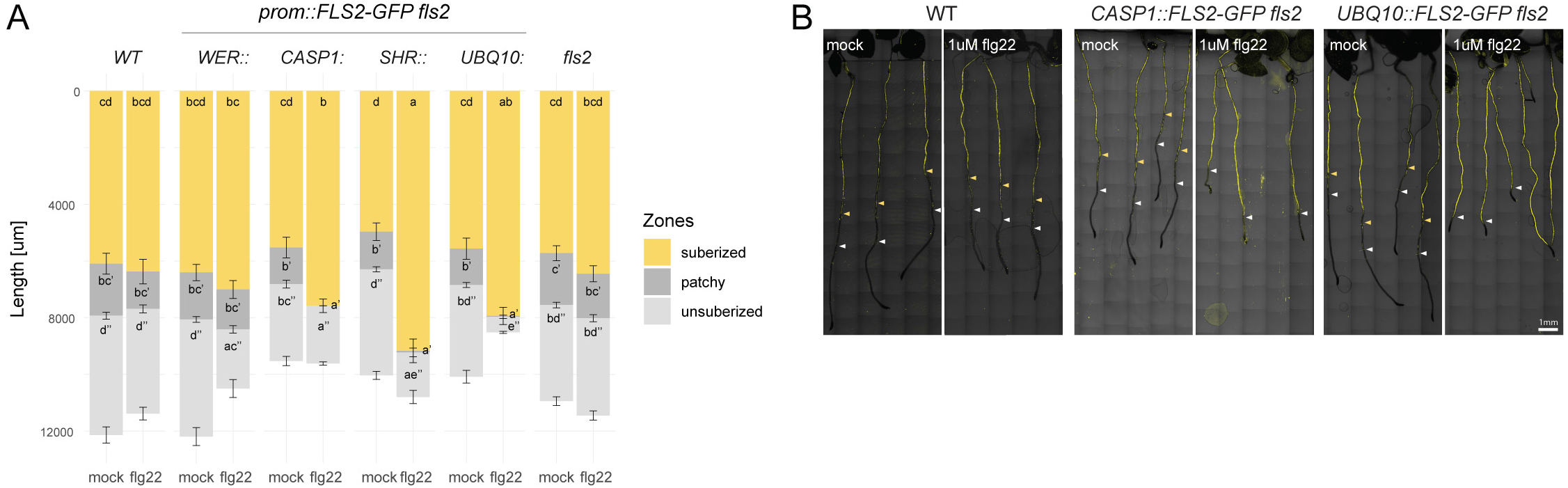
Suberin deposition is triggered by flg22 when endodermal cells expressed *FLS2*. **(A)** Quantification of suberized zone length in seedlings treated for 1 day with 1μM flg22 (18 <= n <= 27). Data of two replicates were pooled. Roots regions were classified as suberized, patchy and unsuberized zones. Error bars represent standard error (SE). Different letters indicate statistically significant differences amongst lines for the specified zone (p<0.05). Multiple comparison was performed using ANOVA and Tukey’s tests for the suberized zone, whereas Kruskal-Wallis and Dunn’s tests were used multiple comparison of patchy and non-suberized zones. **(B)** Whole root views of suberin lamellae deposition in *CASP1::* and *UBQ10::FLS2-GFP fls2* lines compared to wild-type after 1μM flg22 treatment vs mock. Suberin was stained with fluorol yellow. White arrowheads start of patchy zone; yellow arrowheads, start of fully suberized zones. Scale bar, 1mm.

### Super-competent *WER::FLS2* line can detect native bacterial flagellin

The strong impact of flg22 on *WER::FLS2* root growth and cell wall modification prompted us to evaluate whether commensal bacteria would have a similar effects. Indeed, plants that mount ectopic defences in sensitive tissues might suffer from the presence of usually harmless bacteria and tip the balance between growth and defence. The model commensal/beneficial *Pseudomonas protegens* CHA0 does not induce MTI responses in wild-type plants, except at high concentration or if the root is wounded (Zhou *et al*., 2020). However, when inoculated on *WER::FLS2* line, a very evident *PER5* induction could be observed, although no synthetic flg22 peptide was added (Fig.6A). This experiment is therefore a first clear example, where a flg22 response is caused by actual, living bacteria. This flagellin must be released and processed into FLS2-binding smaller peptides (Buscaill *et al*., 2019).

**Figure 6:**
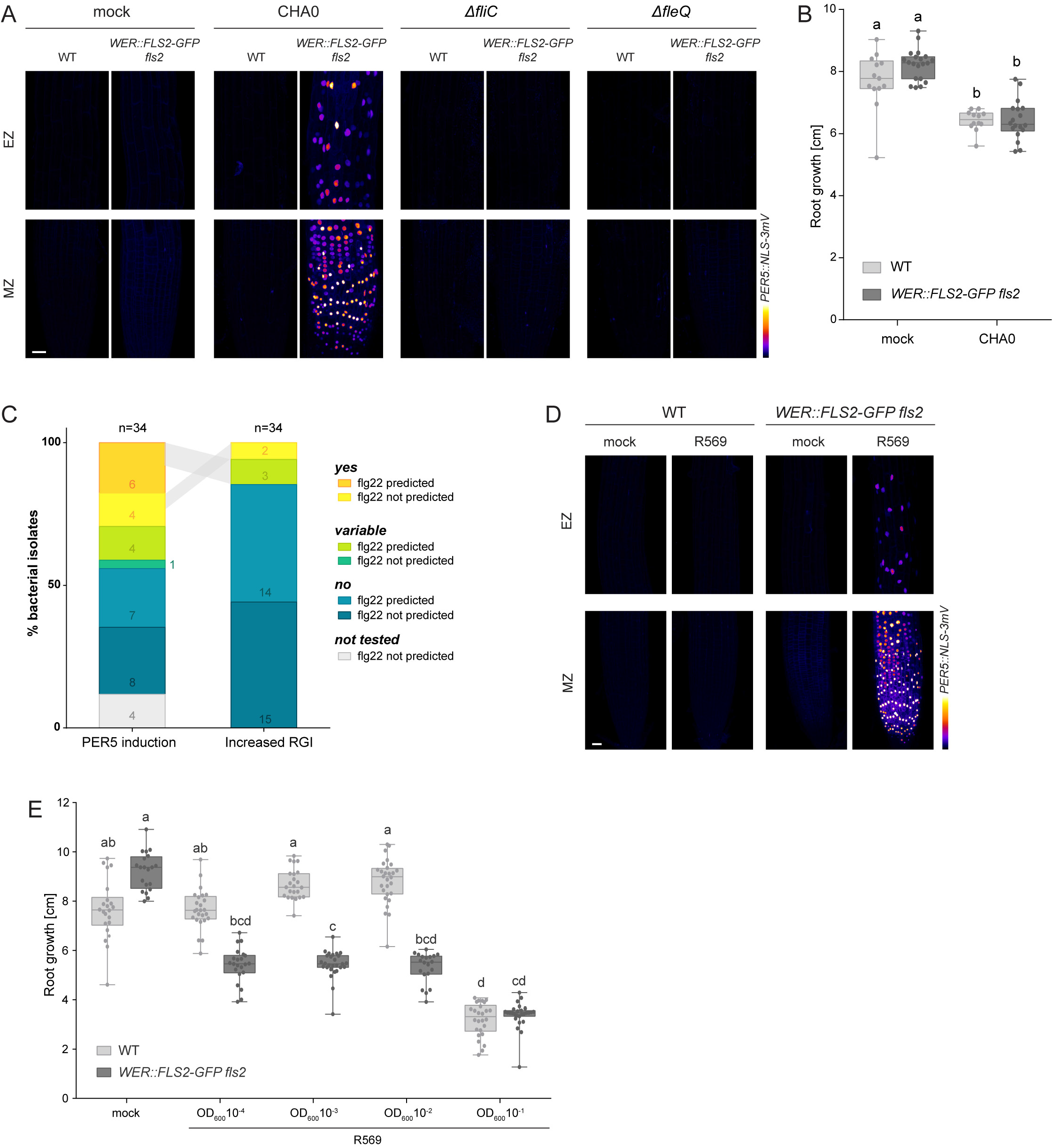
*WER::FLS2* line detects endogenous bacterial flg22. **(A)** CHA0 bacteria trigger a strong induction of *PER5*::*NLS-3mVenus* marker (Fire LUT) on *WER::FLS2-GFP fls2*. Mutants Δ*fliC* and Δ*fleQ* defective for flagellum lose their ability to induce detectable MTI. Δ*fliC* mutant was confirmed by motility assay (see Fig.S7D). Maximum projection of z-stacks imaging meristematic (MZ) and elongation (EZ) zones treated with drop inoculation of bacterial solution of a concentration of OD600 = 0.01 or mock, respectively. Images were acquired at 1dpi. Acquisition done with identical settings. Scale bar, 25μm. **(B)** CHA0 do not induce consistently increased root growth inhibition in *WER::FLS2-GFP fls2*. Root growth was quantified at 6 dpi on plate inoculated with bacteria at OD600 = 10^-3^. Different letters indicate statistically significant differences (p<0.05). Multiple comparison was performed using ANOVA and Tukey’s test. **(C)** Proportion of natural isolates from At-SPHERE culture collection triggering stronger *PER5*::*NLS-3mVenus* induction and increased root growth inhibition (RGI) on *WER::FLS2-GFP fls2* compared to wild-type seedlings (yes), or not (no). Bacteria classified in “variable” presented contradictory results between replicates. Bacteria flg22 sequence was predicted to be recognized by FLS2 (flg22 predicted) or not (flg22 not predicted). Numbers of bacterial isolates in each category are indicated in colour. Grey surfaces indicate identical bacteria strains. **(D)** *Pseudomonas* isolate R569 from At-SPHERE culture collection triggers strong *PER5*::*NLS-3mVenus* (Fire LUT) induction on *WER::FLS2-GFP fls2*. Seedlings were imaged after one-day treatment with OD600 = 0.01. Maximum projection of z-stacks at meristematic zone (MZ) and elongation zone (EZ). Scale bar, 25μm. **(E)** Isolate R569 induces a robust increased root growth inhibition on *WER::FLS2-GFP fls2* compared to wild-type plants. High concentration of bacteria (OD600 = 0.1) is deleterious to both genotypes. Root growth was quantified at 6 dpi on plate inoculated with bacteria at OD600 = 10^-1^ to 10^-4^. Different letters indicate statistically significant differences (p<0.05). Multiple comparison was performed using Kruskal-Wallis and Dunn’s test.

To confirm that the induction of *PER5* was caused by native, bacterial flg22, we infected seedlings with a CHA0 mutant defective for *fleQ*, required for the induction of flagellum development (Arora *et al*., 1997; Kupferschmied, 2015), as well as *fliC*, coding for the flagellin protein (Yamaguchi *et al*., 1984). In contrast to the wild-type strain, Δ*fleQ* and Δ*fliC* mutants could not trigger any response in *WER::FLS2*, demonstrating that defences are induced by the direct FLS2-mediated detection of bacteria-derived flagellin molecules (Fig.6A). We then assessed the impact of CHA0 bacteria on root growth. Surprisingly, despite its induction of PER5, CHA0 did not significantly enhance root growth inhibition in *WER::FLS2* compared to wild-type (Fig.6B). One explanation would be that some commensal bacteria are able to attenuate the excessive MAMP-triggered immune responses in WER::FLS2, thus avoiding root growth inhibition and deleterious defence responses (Garrido-Oter *et al*., 2018; Pel and Pieterse, 2013). Indeed, Ma *et al*. were unable to observe any growth phenotype of *WER::FLS2* plants grown in non-sterile soil. Interestingly, they reported that 41% of root commensals can suppress MAMP-triggered root growth inhibition in mono-associations (*manuscript in preparation*).

### Bacterial community members have diverse impact on *WER::FLS2*

In order to obtain a more comprehensive picture of how *WER::FLS2* affects responses to bacteria, we screened a set of 34 bacterial isolates from the At-SPHERE culture collection of Cologne (Bai *et al*., 2015) for both induction of *PER5* marker and enhanced root growth inhibition in *WER::FLS2* compared to wild-type lines. We selected isolates to represent bacteria from all phyla, with a bias towards bacteria predicted to possess a flg22 peptide sequence recognised by the FLS2 receptor (Fig.6C, Table S1) (Garrido-Oter *et al*., 2018). Amongst the 17 strains predicted to be recognized by FLS2 based on their sequence, only 10 triggered an enhanced *PER5* marker induction in *WER::FLS2*. Moreover, five additional strains, without a predicted recognizable flg22 sequence, had the same effect. This underlines the problematic of predicting flg22 activity from sequence and the potential of the *WER::FLS2* line to rapidly test experimentally, whether a native bacterial flg22 can be detected by the plant.

Although half of bacterial isolates could induce *PER5* marker specifically in the *WER::FLS2* line, only 5 of them affected *WER::FLS2* root growth more strongly than WT, though often with great variation (Fig.6C, Table S1). However, *Pseudomonas* isolate R569 caused strongly enhanced *PER5* induction (Fig.6D) and root growth inhibition compared to WT (Fig.6E). This effect was very robust and was repeatedly observed both in Lausanne and Cologne laboratory growth conditions (Fig.S7C). We demonstrated that commercial, synthetized flg22 from *Pseudomonas aeruginosa* as well as from isolate R569 (flg22R569) similarly induced *PER5* marker expression and inhibited root growth. These effects were abrogated in the *fls2* mutant background (Fig.S7B). We conclude that the commensal R569 isolate induces MTI responses in the *WER::FLS2* line through its native flg22 peptide, which then causes an unbalancing of growth and defence not observed when the bacterium grows on wild-type roots.

## Discussion

It is not understood why only a restricted subset of root tissues can directly respond to MAMPs in the absence of other stimuli (Millet *et al*., 2010; Poncini *et al*., 2017; Zhou *et al*., 2020). The combination of tissue-specific receptor expression and cellular resolution read-outs presented here provides insights into the consequences of altering the spatial patterns of MTI in roots. Our work reveals three important features of MAMP responses.

First, different MTI responses are highly tissue-specific and varying in cell-autonomy. Suberin, for example, is only induced in the endodermis. While *PER5* induction is strictly cell autonomous, *MYB51* and calcium signals are found in cell lacking FLS2 receptor. It will be important to describe larger numbers of response genes for a comprehensive view of MTI. Cell-type specific transcriptomic analysis can complete our understanding of tissue-specific immune pathways (Rich-Griffin *et al*., 2020). Our *prom::FLS2* lines coupled to transcriptional read-outs can now help to distinguish cell-autonomous responses from indirect activation by MTI.

Secondly, we found that the vascular meristem is refractory to flg22 even when expressing FLS2 receptor. The seemingly contradictory finding in Wyrsch *et al*. (2015) can be explained by the whole-organ read-outs used, as well as use of *LBD16::FLS2*, thought to be stele-specific, but that we found to also slightly express in other tissues (Fig.S1J). Lack of downstream signalling components or increased activity of negative regulators could both be responsible for the stele’s inability to respond to flg22. The vascular meristem might be particularly vulnerable to an activation of defence as it contains early-differentiating phloem providing nutrition and hormones to the growing meristem.

Finally, we observed root regions that can be rendered super-competent by FLS2 expression. We speculate that epidermal meristematic cells are not responsive in wild-type (Millet *et al*., 2010; Zhou *et al*., 2020), because only the outer root cap cells can mount MAMP responses that are not detrimental to meristem function. This might be linked to the particular fate of root cap cells that enter apoptosis once they reach the transition zone (Fendrych *et al*., 2014; Kumpf and Nowack, 2015) and excrete mucilage and secondary metabolites influencing root microbiota (Bulgarelli *et al*., 2013; Kumpf and Nowack, 2015). By contrast, epidermal cells might only maintain a competency to respond, if root cap damages by pathogens or other stresses induce *FLS2* expression. Indeed, we clearly showed that constitutive expression of *FLS2* in the meristematic epidermis leads to drastic changes in the root structure upon flg22 treatment in ways that could be detrimental to growth.

Though lignification upon actual bacterial infection is well documented (Lee *et al*., 2019; Nicholson and Hammerschmidt, 1992; Vance *et al*., 1980), treatment with single MAMP was rarely seen to stimulate root lignin deposition (Adams-Phillips *et al*., 2010; Chezem *et al*., 2017; Robertsen, 1986; Smit and Dubery, 1997). Here we show that strong *FLS2* expression reveals the capacity of MTI responses to modify cell walls, probably overriding endogenous negative feedbacks that prevent this from happening in wild-type. This provides an opportunity to study MTI-induced lignification in a simplified and reproducible setting. Interestingly, ectopic corner lignification together with defence genes induction are also observed in response to CIF2 peptide treatment in the endodermis (Alassimone *et al*., 2016; Doblas *et al*., 2017; Fujita *et al*., 2020, 2020; Pfister *et al*., 2014), suggesting the developmental SCHENGEN pathway shares similarities with MTI responses. Nevertheless, lignification is only partly explaining the severe root growth inhibition we observe. Other factors produced in response to flg22 might also interfere with meristem function, such as basic coumarins (Stringlis *et al*., 2019), which inhibit cellulose, resulting in meristem swelling similar to the one observed on *WER::FLS2* (Hara *et al*., 1973).

Our work also reveals that overexpression of a single PRR in a competent, but otherwise non-responsive cell-type, bypasses the absence of visual immune responses to commensal bacteria (Garrido-Oter *et al*., 2018; Millet *et al*., 2010; Yu *et al*., 2019b; Zhou *et al*., 2020). Though bacteria can also inhibit MTI (Couto and Zipfel, 2016; Yu *et al*., 2019a), MAMPs produced by rhizosphere bacteria might often be too low in concentration to activate MTI responses in the first place. Therefore, roots might appear largely unresponsive to bacterial presence without additional stresses (Zhou *et al*., 2020). The obvious root growth phenotype triggered by MTI in *WER::FLS2* lines proves to be a powerful tool to investigate the effect of commensals on root immune responses. Our super-competent lines have allowed for the first time to directly observe stimulation of FLS2 by a native flagellin peptide from an avirulent bacterium. Generally, the cocktail of elicitors that bacteria are thought to release prevent assignment of a MAMP response to an individual MAMP (Tang *et al*., 2017). The *WER::FLS2* line now generates a cell type that responds only to a single MAMP and can test predictions about flg22 peptide detectability, release and processing. Extending our approach, the ectopic overexpression of potential PRR receptors in the epidermal meristem cells could be used to functionally pair novel receptors and ligands.

It has become evident that immune responses cannot be understood without taking into consideration the specificities of different cell type and developmental stages. Our data establishes the necessity for the plant to spatially restrict its immune response. This spatial allocation of defence capacities might in turn influence the microbial colonization pattern of the rhizosphere. The new tools presented will pave the way for a better understanding of bacterial community structures in roots.

## Acknowledgement

We thank the Central Imaging Facility (CIF) of the University of Lausanne for expert technical support. We particularly thank Thomas Boller (Basel), Jean-Pierre Métraux (Fribourg), Silke Lehman (Fribourg) and Ines Wyrsch (Basel) for initiating this project with us. We thank Youssef Belkhadir (Vienna), Thorsten Nürnberger (Tübingen), Corné Pieterse (Utrecht), Cyril Zipfel (Zürich) and Peter Kupferschmied (Bern) as well as all members from the Geldner lab for sharing material and/or helpful discussions and input. Finally, we are grateful to Artan Graf and Yasmine Genolet for their patience and assiduity for the manual curation of cell-type specific quantification. This work was supported by funds to N.G. from an ERC Consolidator Grant (GA-N: 616228–ENDOFUN) and two consecutive SNSF grants (CRSII3_136278 and 31003A_156261). F.Z. was supported by an EMBO Long-Term Fellowship (ALTF 1139-2014).

## Author contributions

A.E. and N.G. conceived, designed, and coordinated the project. A.E., F.Z., J.V., C.M.H., V.D.T. performed all experimental work. A.E. and N.G. wrote the manuscript. A.E., F.Z., J.V., K.M., P.S.L., C.K. and N.G. revised the manuscript and were involved in the discussion of the work.

## Declaration of interests

The authors declare no competing interests.

## Material and methods

### Plant material

*Arabidopsis thaliana* ecotype Columbia Col-0 was used for most experiments. The T-DNA line *FLS2* was obtained from NASC (SALK_062054C) and originally described in (Zipfel *et al*., 2004). The recombinant *WER::FLS2-3myc-GFP*, *CASP1::FLS2-3myc-GFP*, *SHR::FLS2-3myc-GFP*, *UBQ10::FLS2-3myc-GFP, LBD16::FLS2-3myc-GFP* in *fls2* (SAIL691_C04) background, as well as *FLS2::FLS2-3myc-GFP* in Wassilewskija Ws-0 background were provided by Prof. Thomas Boller’s group (Robatzek *et al*., 2006; Wyrsch *et al*., 2015). The defence marker lines *PER5::NLS-3mVenus* and *MYB51::NLS-3mVenus* are described in (Poncini *et al*., 2017). Calcium signalling analysis was carried out thanks to the line *UBQ10::R-GECO1* kindly shared by Prof. Melanie Krebs’s group (Keinath *et al*., 2015).

*PER5*::*NLS-3mVenus* and *MYB51*::*NLS-3mVenus* lines were crossed to the four recombinant lines *WER::, CASP1::, SHR::* and *UBQ10::FLS2-3myc-GFP fls2* lines as well as to the *fls2* mutant. In addition, *UBQ10::R-GECO1* was first crossed to *fls2* mutant, then the resulting homozygous line was crossed again to the four recombinant lines (*WER::/CASP1::/SHR::/UBQ10::FLS2-3myc-GFP fls2*), so that F1 could be directly used for experiments. For quantification of tissue-specific nuclear signal, the constructs *UBQ10::NLS-mTurquoise* or *UBQ10::NLS-tdTomato* were transformed by floral dipping method in all *PER5::/MYB51::NLS-3mVenus* marked *prom::FLS2-3myc-GFP fls2*, *fls2* and wild-type lines (Clough and Bent, 1998). In addition, *RCH1::FLS2-GFP*, *PRP3::FLS2-GFP* and *GRP::FLS2-GFP* were transformed in *fls2* (SALK_062054C).

### Plant growth conditions

For all experiments, seeds were surface-sterilized by gaseous chlorine for 2.5 hours or immersed in a solution of 70% EtOH 0.01% Triton-X-100 for 5 min, washed once in 96% EtOH and dried under sterile conditions. Seeds were stratified in the obscurity for 2 days, then germinated on 1% agar plates containing half-strength Murashige and Skoog (½ MS) medium and 500mg/l MES (Duchefa). Seedlings were grown vertically for 5 days before analysis (otherwise differently specified) in continuous light at 23°C.

For experiments done in Cologne, seeds were sown on 1% Bacto-Agar supplemented with ½ MS with 250mg/l of MES. Seedlings were grown in a light cabinet with short day conditions (10h light-14h dark, 21°/19°C, 65% relative humidity).

### Bacterial strains and growth conditions

*Pseudomonas protegens* strain CHA0 used in this study is a tobacco root isolate with plant-beneficial activities (Stutz *et al*., 1986). CHA0 mutants Δ*fliC* and Δ*fleQ* carrying in-frame deletions in the *fliC* and *fleQ* genes, respectively, were generated using the suicide vector pEMG and the I-SceI system (Kupferschmied, 2015; Martínez-García and Lorenzo, 2011) adapted to *P. protegens* (Kupferschmied *et al*., 2014) with plasmids and primers listed in Supplemental Table S2. The *Pseudomonas* R569 and other natural commensal bacterial isolates were obtained from the At-SPHERE culture collection (Bai *et al*., 2015). CHA0 strains and commensal isolates were routinely cultured at 28°C in, respectively, lysogeny broth (LB) medium (1% tryptone, 0.5% yeast extract and 1% NaCl) or half-strength tryptic soy broth (TSB) (Sigma-Aldrich).

### Plant plasmid construction

Generation of expression constructs was performed with both In-Fusion Advantage PCR Cloning Kit (Clontech) and Gateway Cloning Technology (Invitrogen).

For nuclei labelling with blue fluorochrome, used for quantification, *UBQ10::NLS-mTurquoise2* was generated by triple Gateway recombination reaction using the entry clones pDONR P4-*pUBQ10-*P1R (Zhou *et al*., 2020), pDONR P1-*NLS-mTurquoise2*-P2 and pDONR P2R*-2R3e-nosT*-P3 (Siligato *et al*., 2016) with the destination vector pK7m34GW,0 containing a kanamycin resistance gene for plant selection. For the red version of nuclei labelling, the plasmid *UBQ10::NLS-tdTomato* was used for its FastRed in plantae selection system. Briefly, pDONR P4-*pUBQ10-*P1R (Zhou *et al*., 2020) and pDONR P1-*NLS-tdTomato*-P2 were combined with the destination vector pFR7m24GW by double Gateway reaction. pDONR P1-NLS-tdTomato-P2 was previously generated using in-Fusion cloning to integrate the NLS sequence to pDONR P1-tdTomato-P2.

*RCH1::FLS2-GFP, PRP3::FLS2-GFP* and *GRP::FLS2-GFP* were generated combining the respective entry clones pDONR L4-*pRCH1*-R1 and L4-*pPRP3*-R1 (SWELL lines)(Marquès-Bueno *et al*., 2015), or pDONR L4-*pGRP*-R1 (Andersen *et al*., 2018), with pDon207 containing the L1-*FLS2-3xmyc-GFP*-L2 sequence (Wyrsch *et al*., 2015), in the destination clone pH7m24GW,3.

### Elicitors and inhibitors treatments

Commercial flg22Pa peptide from *Pseudomonas aeruginosa* (QRLSTGSRINSAKDDAAGLQIA) was ordered from EZBioLab. Elf18 oligopeptide from *Escherichia coli* strain GI826 (Ac-SKEKFERTKPHVNVGTIG), *A. thaliana* Plant Elicitor Peptide 1, AtPEP1 (ATKVKAKQRGKEKVSSGRPGQHN) and flg22R569 peptide (NRLSTGKKINSAKDDAAGMQIA) from the isolate *Pseudomonas* R569 were synthesized by Peptide Specialty Laboratories GmbH. (±)-3-Hydroxydecanoic acid (3-OH-C10:0) and chitin were obtained from Sigma-Aldrich. All elicitors were dissolved in deionized MilliQ sterile water at the respective stock concentration of 1mM for flg22Pa, flg22R569, elf18, AtPep1 and 3-OH-C10:0; and 2mg/ml for chitin. For the inhibition of monolignol synthesis, piperonylic acid (PA, Sigma-Aldrich) was dissolved in absolute EtOH at a concentration of 20mM for stock solution.

For elicitor treatments, chemicals were diluted in liquid half strength MS medium (½ MS) to the indicated concentration. Seedlings were grown vertically for 4 days on small ½ MS petri dishes (5.5cm diameter), then 1.5ml of elicitor solution was gently poured over the seedlings to avoid damages induced by transfer. Care was taken that all roots were properly submersed. Seedlings were incubated horizontally for 24h before live imaging analysis of 5-day-old plants or fixation.

For root growth analysis, 5day-old seedlings were carefully transferred on new ½ MS agar plates containing 1μM flg22Pa or flg22R569 and grown vertically for 6 days in standard growth conditions.

For combined treatment with PA and flg22, Col-0 and *WER::FLS2* seedlings were grown for 4 days on ½ MS plates, then were transferred on agar plates supplemented with 10μM PA or ethanol as control. To overcome PA degradation by light but still conserve proper root growth in control conditions, plates were inserted in black boxes open to the top, allowing roots to grow in the obscurity but leaves to reach the light. Roots were hidden from top light using black sterile plastic caches. After overnight pre-treatment, seedlings were again transferred on plates containing 10μM PA/EtOH with/without 1μM flg22/mock, using the same black boxes. Their root tip location was recorded. 48h after the first transfer, root growth was measured and seedlings were fixed as described.

### Microscopy settings and image processing

Imaging was performed on either a Zeiss LSM880, LSM700 or a Leica SP8 inverted confocal scanning microscope. Pictures were taken with a 63x oil immersion objective (Zeiss LSM880), 63x water immersion objective (Leica SP8), 40x water immersion objective (Leica SP8), as well as 20x or 10x dry immersion objectives for tile-scan with 10% overlap (Zeiss LSM880 or LSM700).

The excitation and detection windows were set as following: for visualisation of FLS2 and defence markers, on Leica SP8, GFP/PI (488nm, 500-530nm and 600-670nm); GFP/mVenus/PI (488nm, 490-508nm; 514nm, 517-560nm and 600-670nm, sequential scan), on Zeiss LSM880, GFP/PI(488nm, 500-530nm and 600-650nm respectively), mVenus (514nm, 520-550nm). For lignin analysis: on Zeiss LSM880, calcofluor (405nm, 425-475nm), basic fuchsin (561nm, 600-650nm). For cell-specific quantification: on Zeiss LSM880, DirectRed 23/mVenus/mTurquoise2 (561nm, 580-700nm; 514nm, 520-590nm; 458nm, 460-500nm; sequential scan) and Calcofluor/mVenus/tdTomato (405nm, 415-450nm; 514nm, 520-545nm; 561nm; 570-640nm, sequential scan). For calcium analysis: Zeiss LSM880, R-GECO1 (561nm, 580-640nm). For suberin staining: on Zeiss LSM700, fluorol yellow (488nm, 500-600nm).

Images were processed using the Fiji software. For cross-section maximum projection of MAMP-induced signal (Fig.2B, 2C, S3B, S3C), z-stack pictures were resliced then realigned thanks to the Descriptor-based series registration (2d/3d + t) plugin. A maximum projection of the MAMP marker channel was then merged to a representative single stack of the PI-stained cell wall channel.

### Fixation and staining

Fixation and cell-wall staining were performed according to adapted Clearsee protocol (Kurihara *et al*., 2015; Ursache *et al*., 2018). Briefly, 5-day-old seedlings were fixed for 1h at room temperature under vacuum in 4% paraformaldehyde PBS solution, using 6-well plates, then washed twice for 1min with PBS. Once fixed, seedlings were cleared in Clearsee solution for at least 24h under mild shaking. To visualize cell wall for quantification, clearing solution was exchanged with either 0.1% Calcofluor White or 0.1% Direct Red 23 in Clearsee solution. After at least respectively 30min and 2h of staining, the staining solution was removed and samples rinsed once in fresh Clearsee solution, then washed for 30min in a renewed Clearsee solution with gentle shaking before mounting.

For combined cell wall and lignin staining, fixed and cleared samples were incubated overnight in a Clearsee solution supplemented with 0.2% Basic Fuchsin and 0.1% Calcofluor White. Once the dye solution removed, samples were rinsed once, washed firstly 30min then at least 1.5h before observation.

### Cell-specific quantification

To realize the complete atlas of defence marker responses, the different *prom::FLS2* lines analysed were first transformed with *UBQ10::nls-mTurquoise2* to delimit nuclei. Alternatively, wild-type *PER5::* and *MYB51::NLS-3Venus* lines were transformed with *UBQ10::nls-tdTomato*, which comprise a FastRed rather than a Kanamycin selection. This allowed to quantify directly the T1 and skip one generation. After flg22 treatment, seedlings were fixed in Clearsee and their cell wall stained with DirectRed23, or Calcofluor White respectively. Z-stack were imaged on half section of the roots at 4 different positions, *i.e.* meristematic zone (MZ), elongation zone (EZ), 12 cells and 30 cells after the onset of elongation for 3 to 6 roots by treatment (mock and flg22) and by genotype. Three channels were acquired sequentially for the nuclei (*mTurquoise2* or *tdTomato*), the cell wall (DirectRed23 or Calcofluor White) and the defence markers *PER5* and *MYB51* (mVenus), using the same settings on all pictures for mVenus channel. However, wild-type *UBQ10::NLS-tdTomato* and *prom::FLS2 UBQ10::NLS-mTurquoise2* were imaged with distinct settings due to difference of intensity of the nuclei-labelling constructs. Pictures were processed on FiJi software with a custom batch macro automatizing the following pipeline (Schneider *et al*., 2012). Images were first resliced from the top, then the three channels were separated. A Gaussian blur was applied on the nuclear and cell wall marker channels, while the PTI marker channel was left untouched to not affect the signal to measure. In a second step, the cell wall channel was subtracted to the nuclear channel to reduce the unspecific background noise of the *UBQ10::nls-mTurquoise2* marker. The “cleaned” nuclear marker channel was transform to 8 bits to facilitate further processing.

We then used the 3D suite to generate a 3-dimensional Region Of Interest (ROI) for each nucleus (Ollion *et al*., 2013). We first applied the plugin 3D iterative thresholding on the 8bits-cleaned-nuclear marker channel (Gul-Mohammed *et al*., 2014). In this process, all possible thresholds are tested, which will detect objects for all thresholds. Subsequently, the algorithm will define the best object segmentation for each of the object, which means that different objects can be segmented with different threshold. This is particularly useful to detect objects with variable intensity in an uneven background, to which a single intensity threshold would either miss many objects or include background noise. We used the following settings: min vol pix = 250, max vol pix = 10000, min threshold = 0, min contrast (exp) = 5, criteria method = COMPACTNESS, threshold method = STEP, Segment results = All, value method = 10.0, Starts at mean = on. The plugin gives as output the 3D threshold delimiting all the future ROIs, *i.e.* the nuclei to quantify. It must be noted that depending on the pictures, some nuclei can be missed, or false positive can be added, but all pictures were then manually curated in a later step. The output came as 2-channels-images, whose last channel is completely black and can be removed by the splitting channel function.

We then use the 3D object counter plugin to define all ROIs, based on the 3D threshold obtained previously, and to redirect the analysis on the defence marker channel (Bolte and Cordelières, 2006). Options were set using the 3D OC Options as following: all parameters were selected, *i.e.* “Volume”, “Nb of Obj. voxels”, “Nb of Surf. voxels “, “Integrated Density”, “Mean Gray Value”, “Std Dev Gray Value”, “Median Gray Value”, “Minimum Gray Value”, “Maximum Gray Value”, “Centroid” “Mean distance to surface”, “Std Dev distance to surface”, “Median distance to surface”, “Centre of mass”, and “Bounding box”. In addition, we ticked both parameters “Close original images while processing” and “Show masked image. The maps’ parameters were set as follows: dots size = 5, font size = 12, “Show numbers” and “White numbers” were ticked. Importantly, the “Results Table Parameters” should be set on: “Store results within a table named after the image”, which allows to keep track of the files in batch mode. Finally, the measures were “Redirected to” the defence marker channel. After setting all the parameters, the analyse “3D Object Counter” was run. Threshold was set to 1 and minimum size filter to 10. The following maps and result tables were asked to be shown: objects, centroids, statistics, summary.

The process gives in output four different files. The “Centroid map” shows the centre of each ROI by a dot, numbered accordingly. The “Object map” is the representation of all ROIs, each of them being numbered. Our macro merge this map to both the cell wall marker channel and the nuclei channel for later identification of nuclei. The 3D Object counter also provides a .csv file called “Statistics” with all parameters measured for each nucleus. The last file provided is the “Masked image”, which represent the defence marker channel masked by the threshold.

To terminate the quantification, each data set was manually curated to assign a tissue (epidermis, cortex, endodermis, stele or root cap) to each nucleus. A maximum of around 20 nuclei by tissue type and by picture were identified. “Mean Gray Values” was extracted and use for analysis. Mean nuclear intensity for each genotype, treatment, root region and tissue were calculated and colour coded using the heatmaply() function in R (heatmaply library). Atlas maps were drawn according to these coloured values. Fold changes for *MYB51* induction were calculated and colour-coded using the same procedure.

### Ca^2+^ imaging on roots and quantification

For calcium responses analysis, *UBQ10::R-GECO1* samples were mounted as follows. Seedlings, once at a time, were glued to a large (60mm) coverslip previously sprayed with medical adhesive (Adapt Medical Adhesive Spray, Hollister). A silicon isolator (Grace Bio-Labs Press-to-seal silicon isolator, No PSA, 20mm diameter, Sigma) was then quickly placed around the seedling and 600µl of sterile germination medium (0.75mM CaCl2, 1mM KCl, 0.25mM Ca(NO3)2•4H20, 1mM MgSO4•7H20, 0.2mM KH2PO4, 50μM NaFe(III)EDTA, 50μM H3BO3, 5μM MnCl2•4H20, 10μM ZnSO4•7H20, 0.5μM CuSO4•5H20, 0.1μM Na2MoO3, pH adjusted to 5.6 with NaOH) was dropped on the root. The drop was spread with a pipet tip to cover the whole surface delimited by the silicon isolator and the seedling let to rest for at least 20min. For full root imaging, tile scans combined to time laps were performed under Zeiss LSM880 confocal laser scanning microscope with 20x objective as described above. As few tiles as possible were selected to limit time acquisition, no averaging was done, and pinhole was entirely open. Images were taken continuously, with an average time interval of 5 to 7 seconds. Acquisition of baseline signal was performed for 5min, then 7.5μl of 100μM flg22 diluted in water was added to the germination medium solution. Acquisition was continued for at least 20min. For tissue-specific imaging and quantification, small z-stack (∼ 8 slices) with 5μm intervals were taken on half a root in the elongation zone for wild-type and *WER::FLS2* samples, or in the differentiated zone for *CASP1::/SHR::/UBQ10::FLS2*.

R-GECO1 signal was quantified for each tissue on the z-stack acquisition. ROIs delimiting a tissue type were drawn manually on the most appropriate stack (*i.e.* that presents a clear surface view if possible), using maximum projection of 2 stacks when necessary.

Fractional fluorescence changes ΔF/F were calculated for each ROI from background corrected intensity values as (F-F0)/F0, where F0 is equal to the average fluorescent intensity of the baseline of the measure, on 4 min from t=0.

### Suberin staining

To highlight suberin lamellae, seedlings were fixed and stained with the methanol-based fluorol yellow staining protocol as described in (Fujita *et al*., 2020). Samples were imaged using the Zeiss LSM700 as described above.

### Bacterial root inoculation assays

PTI assays were performed by drop inoculation on agar plates. Briefly, 2μl of bacterial suspension (cells centrifuged and resuspended in fresh LB or 50% TSB for CHA0 and R569, respectively) of OD600 = 0.01 was added to the tip of 5-day-old seedlings. Once the drop dried, seedlings were grown vertically in standard conditions for 1 to 3 days. For the fast screening of bacterial isolates, roots were observed under a Leica DM 5500B epifluorescence microscope (GFP lamp). Representative pictures of roots were imaged using confocal scanning microscopy (Leica SP8) after a short wash in deionized H2O.

Root growth inhibition assays were completed on agar plates inoculated with bacteria at mentioned concentrations. Briefly, bacterial cultures were grown as previously described in 50% TSB, then centrifuged and resuspended in fresh medium. OD600 was measured and adjusted to 100x the desired concentration. 500μl of concentrated bacterial inoculum was then added to 50ml of semi solid ½ MS medium afore cooled down to around 30°C. Inoculated media were gently mixed by inverting several times, then poured in square petri dishes. Five-day-old *WER::FLS2* and wild-type Col-0 seedlings previously grown on mesh (15mm x 100mm, on top of the plate), were transferred with sterile forceps on the inoculated plates. Seedlings were selected for similar root size, the ones being obviously too long or too short removed from the mesh with sterile toothpicks. After transfer, root tip locations were marked for keeping track of growth, then plants were grown in standard conditions for 6 days. One day post-inoculation, root tip positions were again recorded, and all seedlings that completely stopped to grow were dismissed from the analysis. This ensured that only seedlings that recovered properly from the transfers were considered. Plates were scanned at 6 dpi and root growth measured using Fiji plugins “Simple Neurite Tracer” (Frangi *et al*., 1998).

### Statistical analysis

Statistical analyses were done using R3.6.0 or Graphpad Prism 7.0 software (https://www.graphpad.com/). For multiple comparison, ANOVA followed by Tukey’s Honestly Significant difference (HSD) test were applied when linear model assumptions were met. On the contrary, Kruskal-Wallis test followed by Dunn’s multiple comparison test were performed. For analysis of suberization along the roots, comparisons were performed for each zone separately, and different letters indicates significant differences for a given zone (a, b, c or a’, b’, c’ or a’’, b’’, c’’).

## Supplemental Information Titles and Legends

**Supplemental Figure S1:**
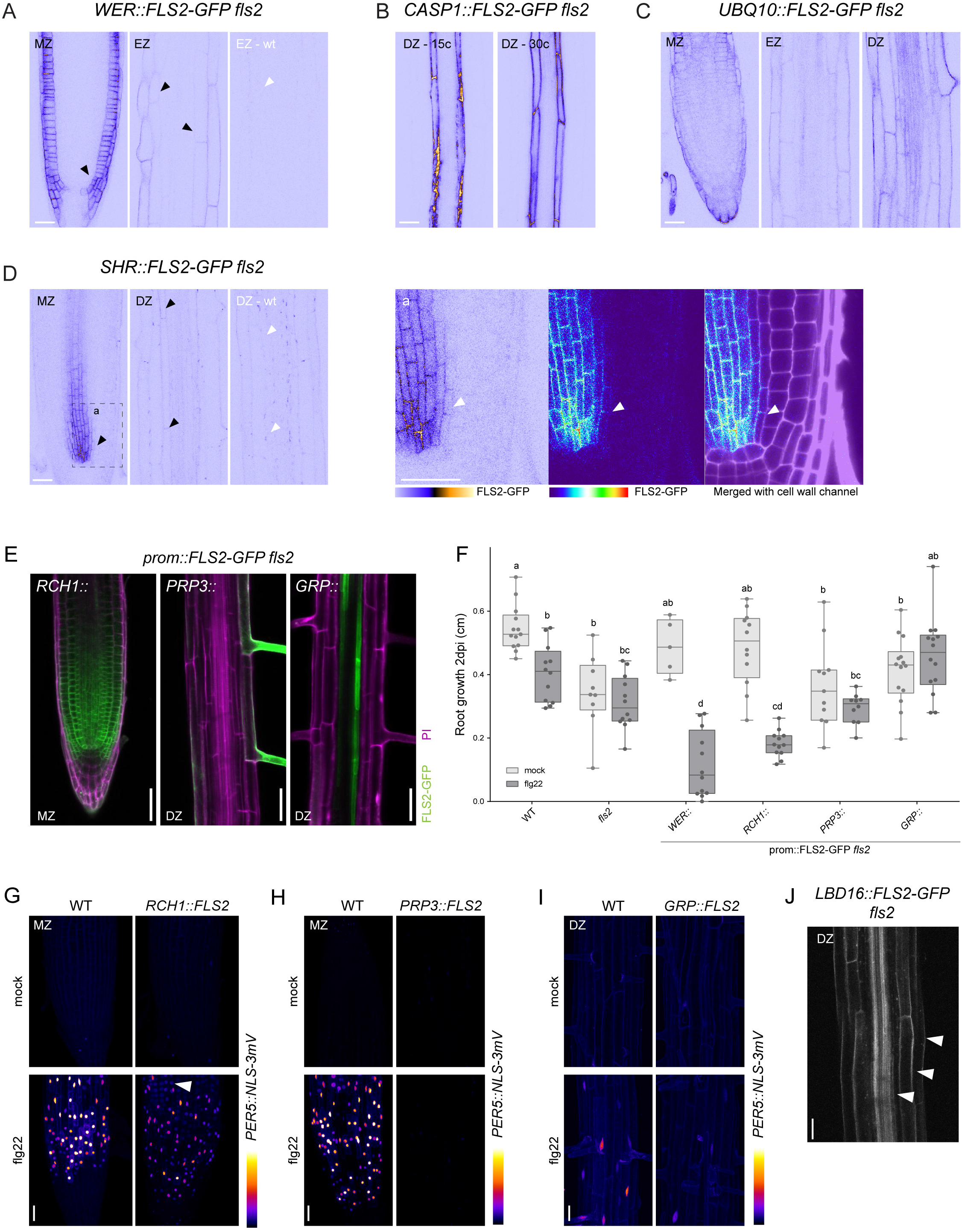
Expression pattern of *prom::FLS2* complementing *fls2*. **(A)** *WER::FLS2-GFP* expression. *WER* promoter expressed principally *FLS2* in epidermal cells, but some weak signal can be observed in cortex (black arrowheads). Picture of wild-type plants taken with identical setting (EZ-wt) is showed for comparison (cortical cell, white arrowhead). **(B)** *CASP1::FLS2-GFP* is expressed exclusively in endodermal cell line in early and later differentiated zones (15 cells respectively 30 cells after onset of elongation). **(C)** *UBQ10::FLS2-GFP* is expressed in all tissue types in every region of the root. **(D)** *SHR::FLS2-GFP* is expressed strongly in the stele of the meristem then decreases in intensity in later regions. Some weak signal can be detected in endodermal cells (black arrowheads). Picture of wild-type plants taken with identical setting (DZ-wt) is shown for comparison (endodermal cells, white arrowheads). Close-up view of dashed squared box is found in (a). FLS2-GFP (visualized by ICA and Thermal LUTs) is merged with cell wall stained by PI (white). White arrowheads point at endodermal cells expressing weakly *FLS2*. **(E)** *RCH1* promoter expresses *FLS2* in the meristem, *PRP3::* in the root hair cells and *GRP::* in the pericycle cells. FLS2-GFP (green) is co-visualized with PI-stained cell wall (magenta). **(F)** Flg22 treatment increases root growth inhibition in *WER::* and *RCH1::FLS2-GFP fls2* hypersensitive line only. Root length quantification of *prom::FLS2-GFP fls2* lines treated with 1μM flg22 for 2 days. Boxplot centre represents the median (5 <= n <= 14). Different letters indicate statistically significant difference between means by 2-ways ANOVA and Tukey’s multiple comparison. **(G)** Maximal projection *PER5::NLS-3mVenus* marker (Fire LUT) in *RCH1::FLS2-GFP fls2* compared to WT shown for MZ. Seedlings were treated for 24h with 1μM flg22. Images were taken with identical settings. White arrow, epidermal signal. **(H)** Maximal projection *PER5*::*NLS-3mVenus* marker (Fire LUT) in *PRP3::FLS2-GFP fls2* compared to WT shown for MZ. Seedlings were treated for 24h with 1μM flg22. Images were taken with identical settings. **(I)** Maximal projection *PER5::NLS-3mVenus* marker (Fire LUT) in *GRP::FLS2-GFP fls2* compared to WT shown for the DZ. Seedlings were treated for 24h with 1μM flg22. Images were taken with identical settings. **(J)** *LBD16* promoter expresses *FLS2-GFP* in all tissues in the differentiated zone (DZ). Note that in contrast to previous report, FLS2 is present in epidermis, cortex and endodermis (white arrows) in addition to the stele. Meristematic zone (MZ), elongation zone (EZ), differentiation zone (DZ). Scale bar, 25μM.

**Supplemental Figure S2:**
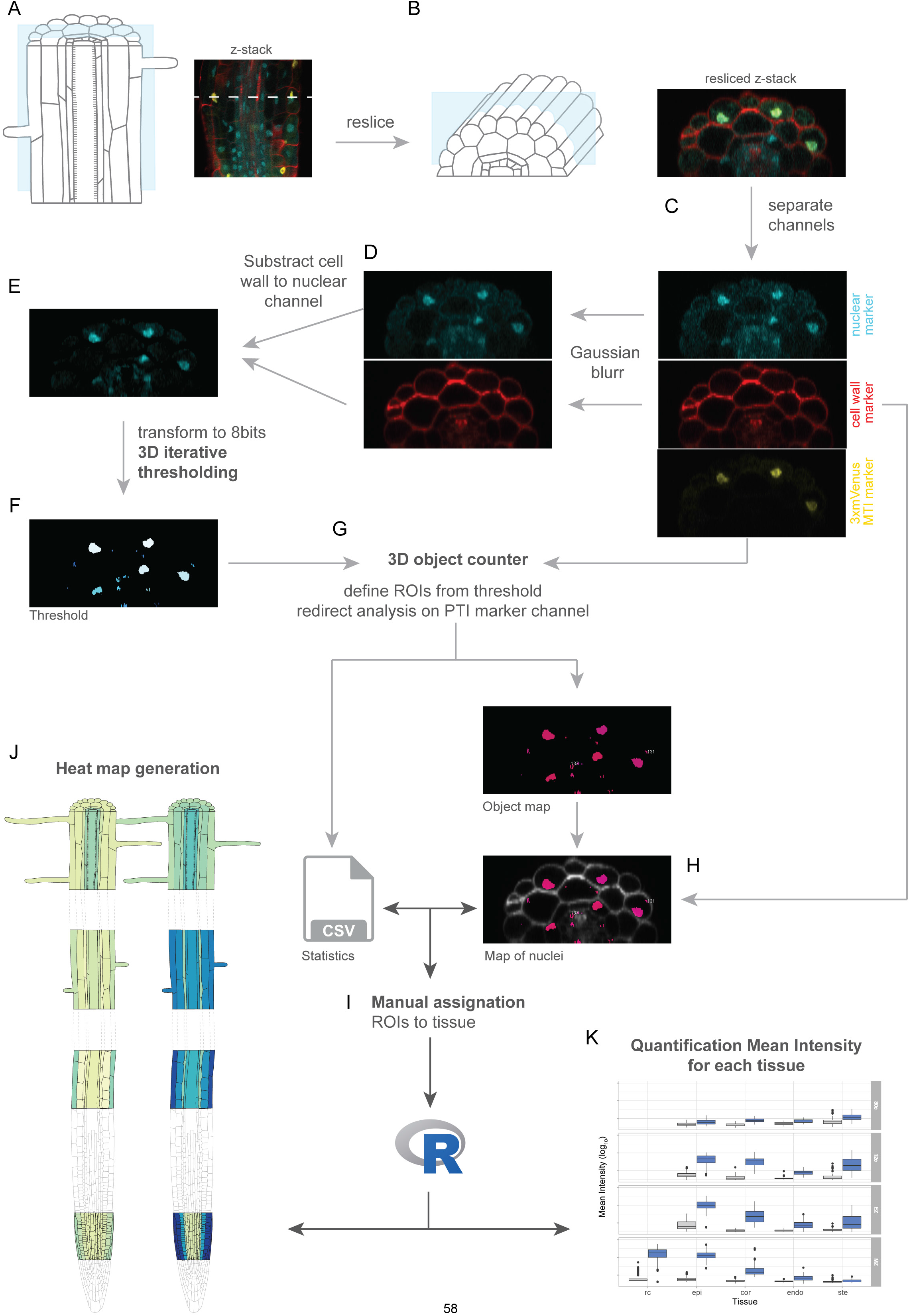
Quantification procedure. **(A)** Z-stack images with 3 channels (red: cell wall, blue: *UBQ10::NLS-mTurquoise/tdTomato*, yellow: *PER5*/*MYB51*::*NLS-3mVenus*) were taken from 4 different regions of the root (meristematic zone, elongation zone, 12 cells and 30 cells after onset of elongation), for 3-6 seedlings by treatment by genotype. **(B)** Each Z-stacks are resliced to get cross-section view. The three channels are separated **(C)** and a Gaussian blur filter is applied on the cell wall and the MTI marker channel **(D).** Blurred cell wall channel is then subtracted from blurred nuclear marker channel to remove non-nuclear background **(E)**. The obtained cleaned nuclear channel is then converted to 8-bit and a 3D iterative thresholding is performed to delimit ROI for each nuclei **(F).** The 3D object counter plugin is then used to measure the mean signal intensity of each nuclei delimited by the obtained ROIs in the MTI marker channel. The plugin gives as output a .csv file with the measured values, a masked image of the PTI marker channel and an object map, delimiting the identified nuclei **(G)**. The object map is then coupled to the original cell wall marker to define the tissue origin of each nuclei **(H)**. Each map was then reviewed manually to assign 20 nuclei for each cell type and to complete .csv files **(I)**. Average of the mean signal intensity of each nuclear tissue-specific signal were calculated, transformed into log10 and colour coded using the heatmaply() function in R **(J)**. Boxplots were generated to represent signal variability **(K)**.

**Supplemental Figure S3:**
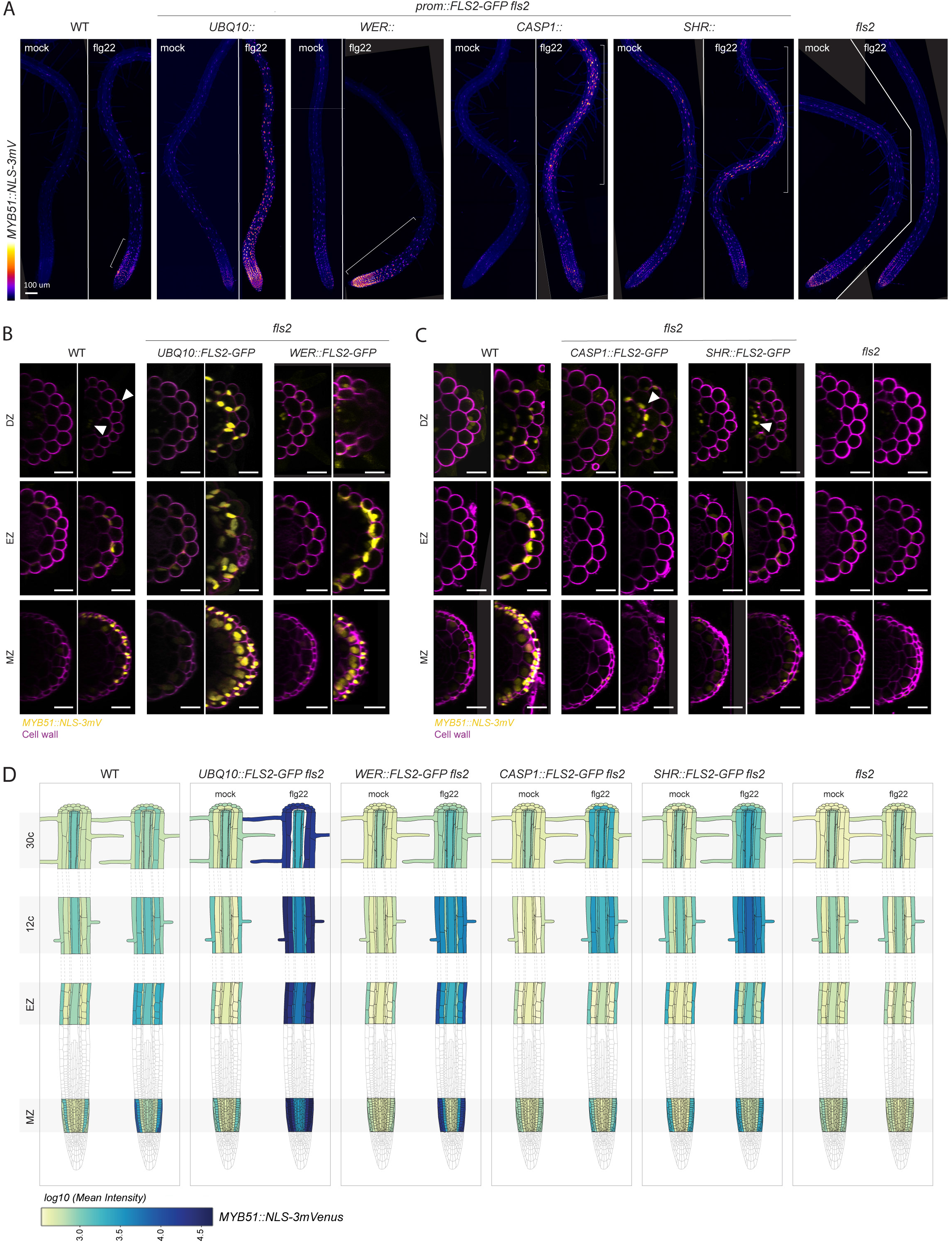
*MYB51* marker is induced non cell-autonomously by flg22 treatment. **(A)** Overview of *MYB51*::*NLS-3mVenus* response to 1μM flg22 after 1 day in different *prom::FLS2-GFP fls2* lines. *MYB51* zone of responsiveness follows *FLS2* expression pattern. Tile scan images were taken with similar settings. Settings are always identical between mock and corresponding flg22 treatment. Brackets indicate zone of responsiveness. Scale bar, 100μM. **(B)** Maximal projection of transverse sections views of *MYB51* expression pattern in UBQ10:: and *WER::FLS2-GFP fls2* compared to WT shown for meristematic zone (MZ), elongation zone (EZ) and differentiated zone (DZ, 30 cells after start of elongation). Seedlings were treated for 24h with 1μM flg22. Nuclear localized mVenus signal (yellow) was co-displayed with propidium iodide cell wall marker (PI, purple). Images were taken with similar settings, while corresponding mock and flg22 treatment pictures for each zone separately have identical parameters. Pictures were acquired with low gain compare to Fig.S2C due to strong average intensity of UBQ10:: and *WER::FLS2-GFP fls2* responses, explaining the faint signal in WT (white arrowheads). Scale bar, 25μm. **(C)** Maximal projection of transverse sections views of *MYB51*::*NLS-3mVenus* expression in *CASP1::* and *SHR::FLS2* as well as WT and *fls2*. *MYB51* expression pattern stay conserved (epidermis-cortex-stele), but intensity is increased in neighbourhood of cells expressing *FLS2*, such as in cortex in *CASP1::FLS2-GFP fls2* or stele in *SHR::FLS2-GFP fls2* (white arrowheads). Imaged were acquired as Fig.S2B., with similar settings between genotypes, while corresponding mock and flg22 treatment pictures have identical parameters. Due to lower average signal intensity, pictures were acquired with increased gain compare to Fig.2B. Scale bar, 25μM. **(D)** Quantitative map of *MYB51*::*NLS-3mVenus* responses inferred from tissue-specific quantification after 24h treatment with 1μM flg22. Nuclear signals were quantified in ROI delimited with *UBQ10::NLS-mTurquoises2* for all tissue-specific promoter lines, while wild-type (WT) signal was quantified with *UBQ10::NLS-tdTomato* marker. Mean intensity is comparable between *prom::FLS2-GFP fls2* lines but not to wild-type. Note the constitutive signal present in untreated seedlings.

**Supplemental Figure S4:**
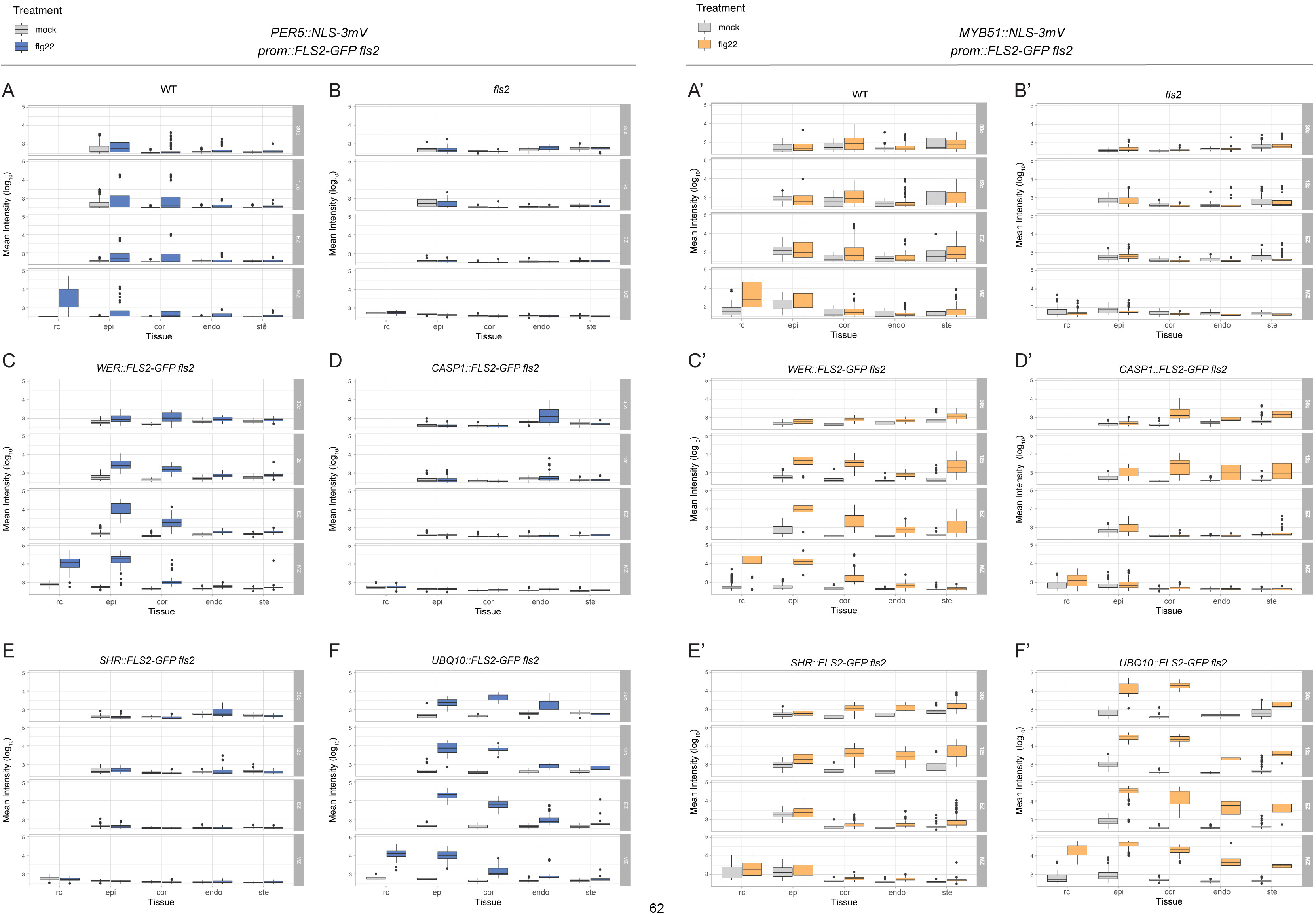
*PER5* and *MYB51* tissue-specific quantification values. Boxplots for mean intensity of *PER5*::*NLS-3mVenus* **(A-F)** and *MYB51*::*NLS-3mVenus* **(A’-F’)** marker calculated from tissue-specific nuclear signals for **(A)** wild-type plants, **(B)** *fls2* mutant, **(C)** *WER::FLS2-GFP fls2*, **(D)** *CASP1::FLS2-GFP fls2*, **(E)** *SHR::FLS2-GFP fls2* and **(F)** *UBQ10::FLS2-GFP fls2*. Boxplot centre represents the median. MZ, meristematic zone; EZ, elongation zone; 15c, 15 cells after onset of elongation; 30c, 30 cells after onset of elongation; rc, root cap; epi, epidermis; cor, cortex; endo, endodermis; ste, stele.

**Supplemental Figure S5:**
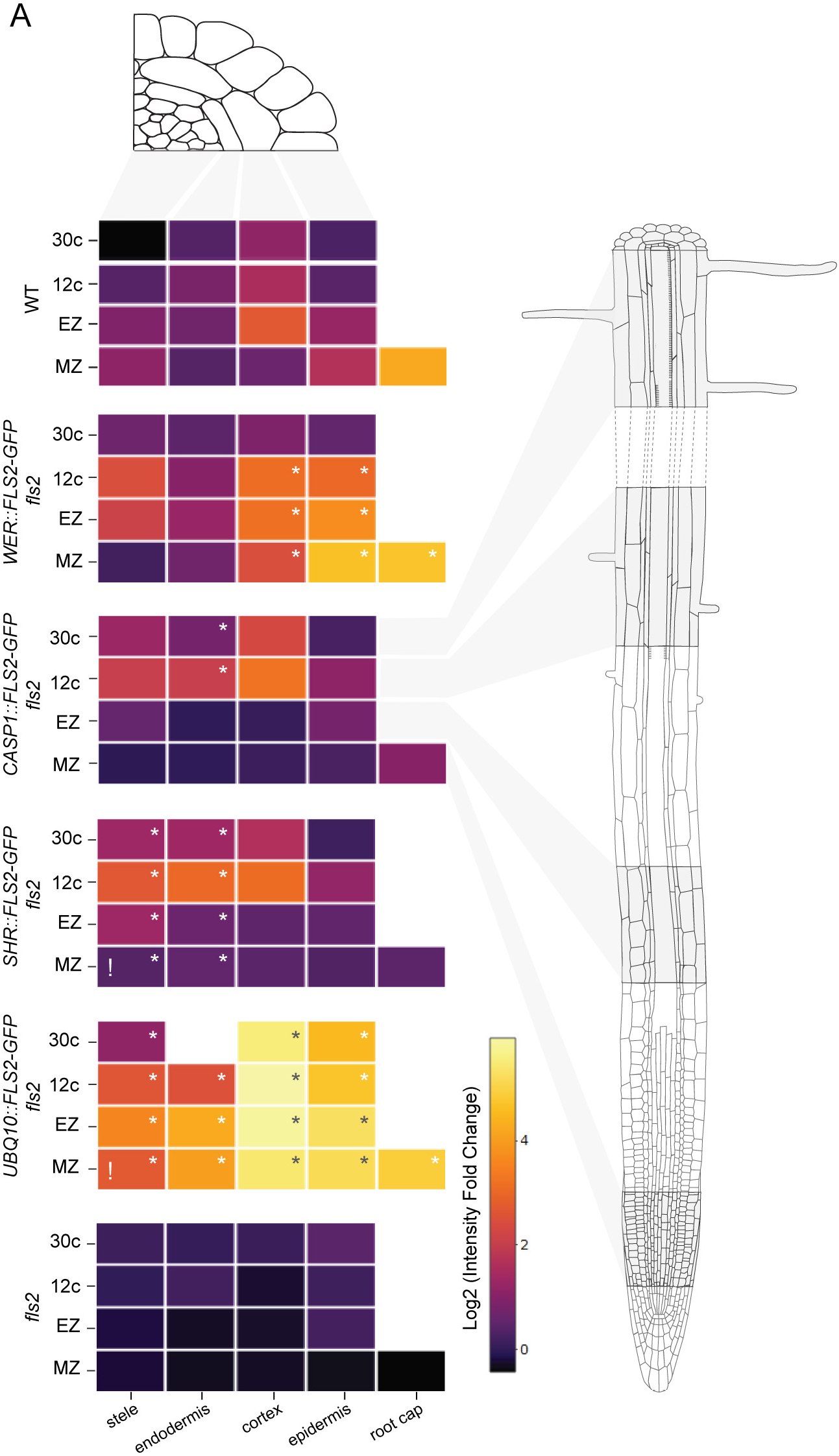
Tissue-specific quantification of*MYB51* fold change. Log2 transformed fold change of intensity of *MYB51::NLS-3mVenus* in WT, *fls2* and the different *prom::FLS2-GFP fls2* lines. Pattern of induction of *MYB51* changed between the different lines but increased signal is not restricted to tissue expressing *FLS2* (stars). Note that *MYB51* can be induced in the stellar meristem in *UBQ10::FLS2* but not in *SHR::FLS2* (!). MZ, meristematic zone; EZ, elongation zone; 15c, 15 cells after onset of elongation; 30c, 30 cells after onset of elongation; rc, root cap; epi, epidermis; cor, cortex; endo, endodermis; ste, stele.

**Supplemental Figure S6:**
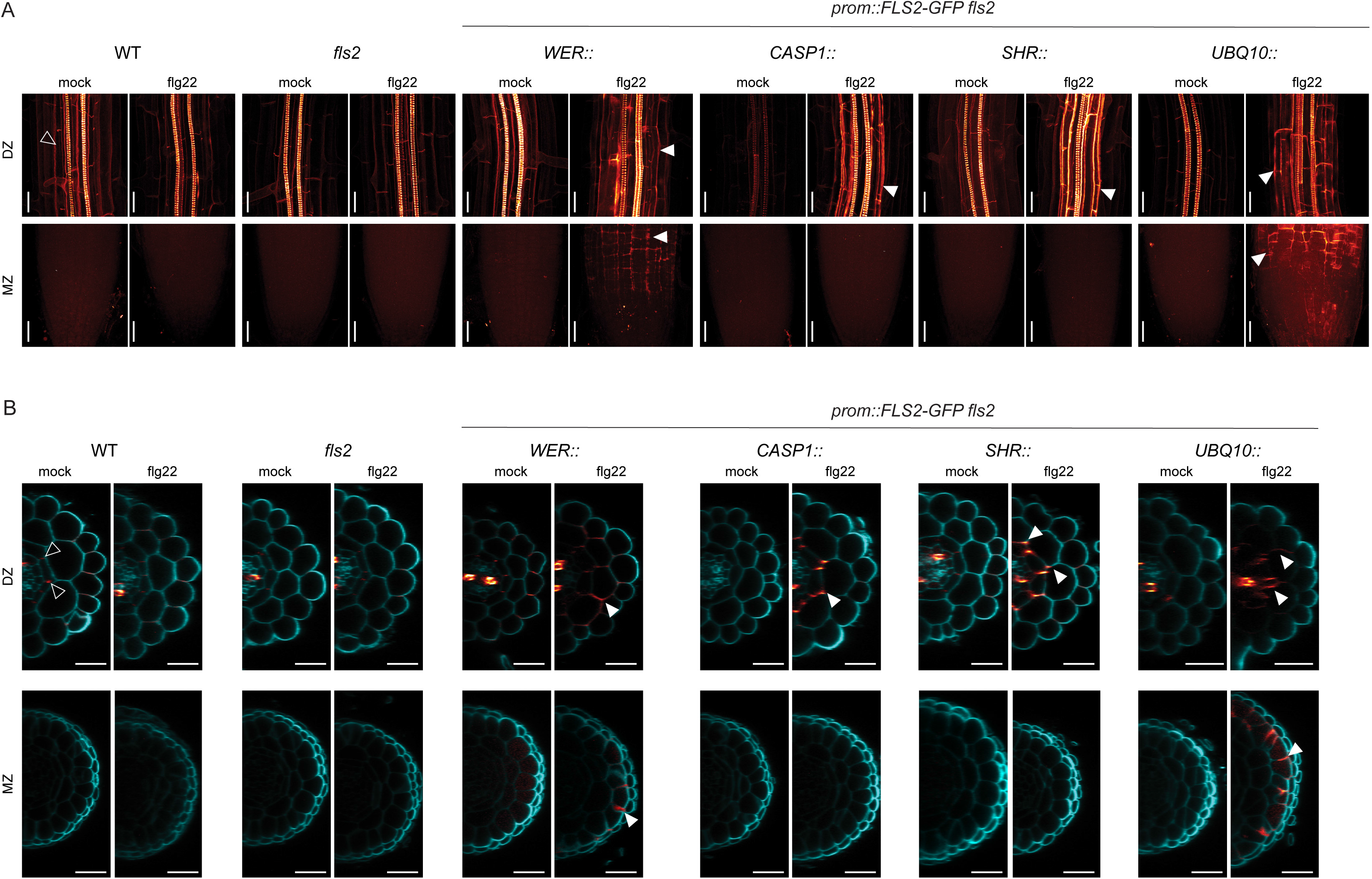
Lignin deposition is a cell-autonomous process. **(A)** Maximum projection showing lignin deposition stained by basic fuchsin in the meristematic zone (MZ) and the differentiated zone (DZ) of the different *prom::FLS2-GFP fls2* lines after 1 day treatment with 1μM flg22. While neither wild-type nor *fls2* roots show lignin deposition outside of the xylem and the endodermal Casparian strip barrier, WER:: and *UBQ10::FLS2-GFP fls2* lines deposit lignin in both MZ and DZ. In contrast, CASP1:: and *SHR::FLS2-GFP fls2* lignified heavily the DZ only. Black arrowheads, Casparian strip. White arrowheads, ectopic lignin deposition. Scale bar, 25μM. **(B)** Cross section of z-stack presented in (A). Cell wall stained with calcofluor (blue) is co-visualized with lignin stained with basic fuchsin (red). *WER::FLS2-GFP* expression drives lignin deposition between cortex and epidermal cells in DZ, and between epidermal cells and root cap in MZ. This pattern is also observed in *UBQ10::FLS2*, but extends to cortex and endodermis in DZ. Both *CAPS1::* and *SHR::* deposit lignin ectopically between cortex and endodermal cells after flg22 treatment. White arrowheads, ectopic lignin. Black arrowheads, Casparian strip. Scale bar, 20μM.

**Supplemental Figure S7:**
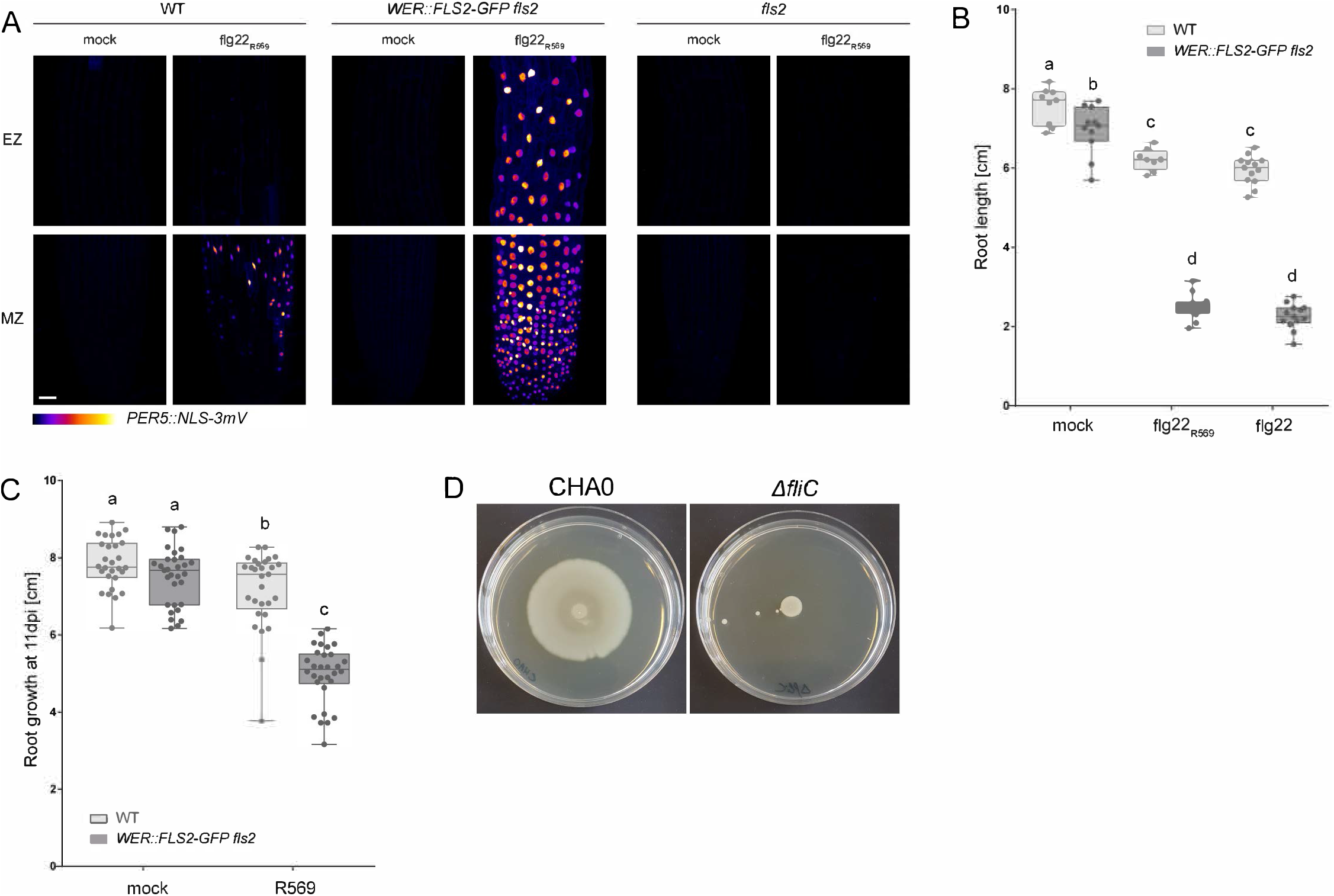
Flg22 from bacterial isolate *Pseudomonas* R569 is recognized by *WER::FLS2*. **(A)** flg22R569 triggers a strong induction of *PER5*::*NLS-3mVenus* marker (Fire LUT) on *WER::FLS2-GFP fls2* compared to wild-type plant, but the detection is abolished in the *fls2* mutant. Maximum projection of z-stacks imaging meristematic (MZ) and elongation (EZ) zones treated for 1 day with 1uM flg22R569. Acquisition done with identical settings. Scale bar, 25μm. **(B)** flg22R569 inhibits root growth weakly on wild-type (WT) and strongly on *WER::FLS2-GFP fls2* in the same extent than commercial flg22 for *P. aeruginosa*. Seedlings were transferred for 7 days on plates containing 1uM flg22, flg22R569 or mock. Boxplot centre represents the median. Different letters indicate statistically significant difference (p<0.05) between means by 2-ways ANOVA and Tukey’s multiple comparison tests. **(C)** Bacterial isolate R569 induces stronger root growth inhibition on wild-type seedlings (WT) than on *WER::FLS2-GFP fls2*. Replicate carried out in Cologne with different growth conditions (see material and methods). Five-days old seedlings were transferred for 11 days on plate containing bacteria at a concentration of OD600 = 0.01. Boxplot centre represents the median. Different letters indicate statistically significant difference (p<0.05) between means by ANOVA and Tukey’s multiple comparison tests. **(D)** Motility assay for CHA0 and its Δ*fliC* mutant.

### Supplemental Tables

**Supplemental Table S1:**
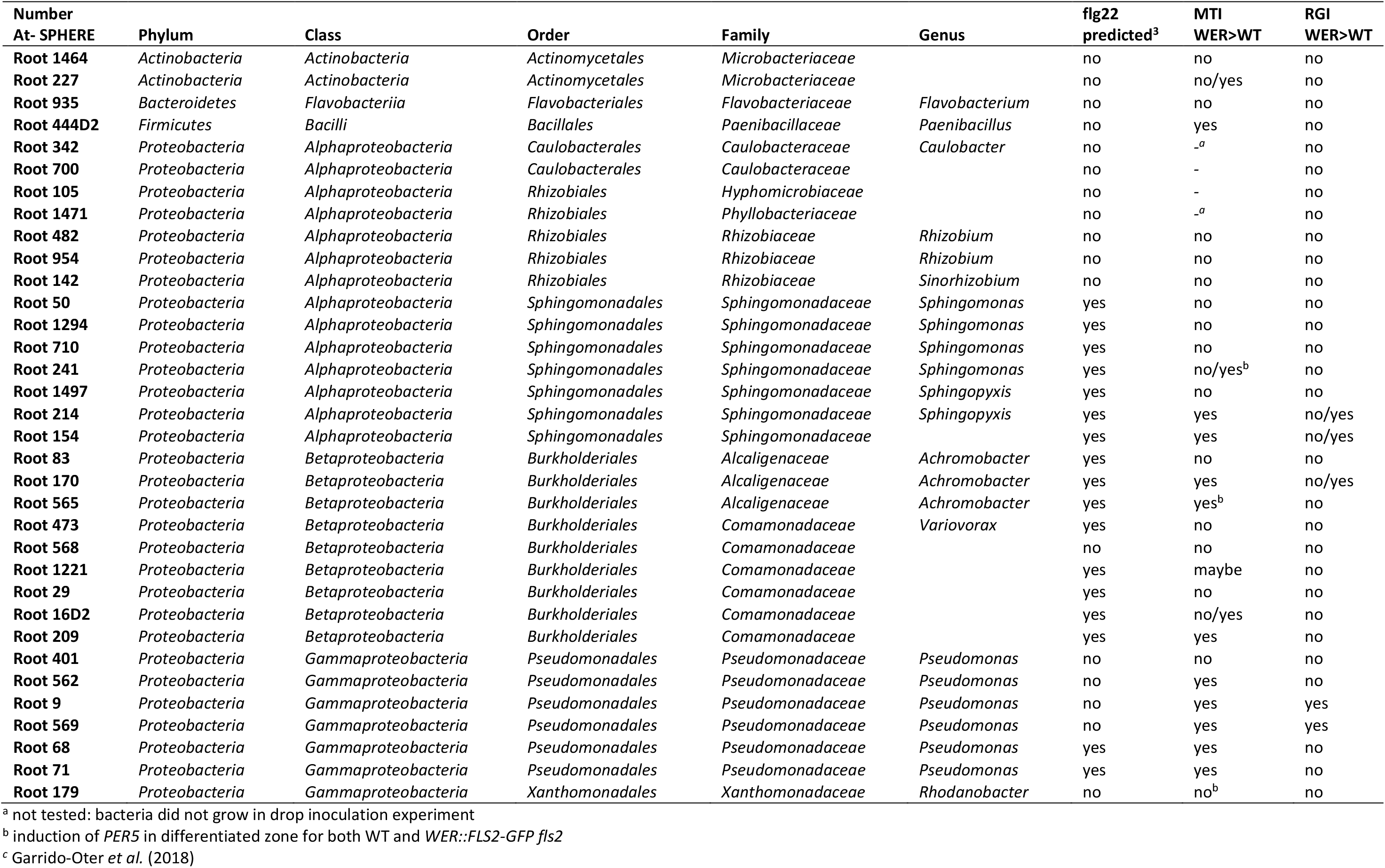
Summary of bacteria screen for PTI assay and RGI assay

**Supplemental Table S2:**
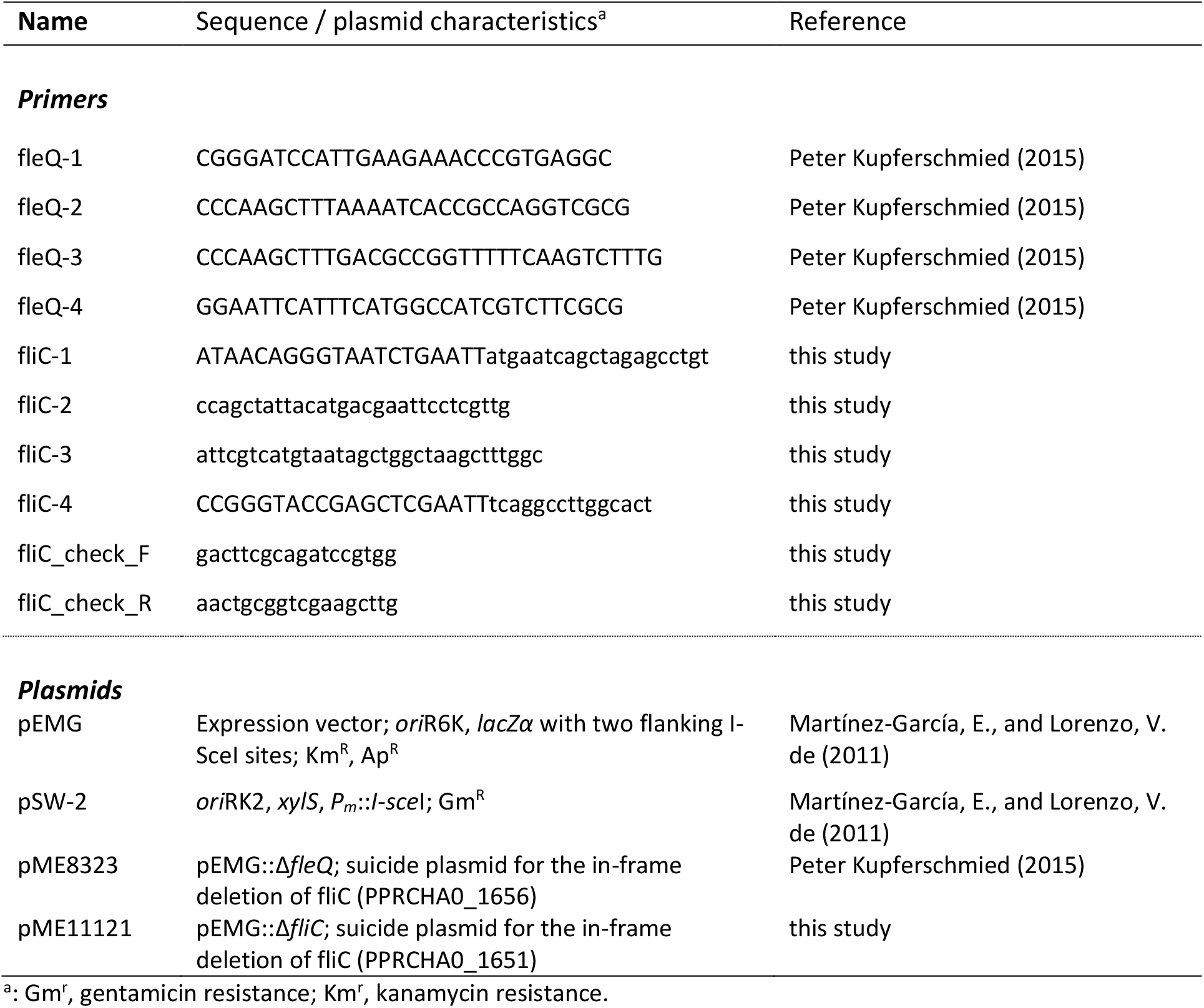
Primers and plasmids used for bacterial mutagenesis.

### Supplemental Videos Titles

Movie 1: Treatment of *UBQ10::R-GECO1* in WT background with 1.25uM flg22 – overview

Movie 2: Treatment of *UBQ10::R-GECO1 WER::FLS2-GFP fls2* with 1.25uM flg22 – overview

Movie 3: Treatment of *UBQ10::R-GECO1 CASP1::FLS2-GFP fls2* with 1.25uM flg22 – overview

Movie 4: Treatment of *UBQ10::R-GECO1 SHR::FLS2-GFP fls2* with 1.25uM flg22 – overview

Movie 5: Treatment of *UBQ10::R-GECO1 UBQ10::FLS2-GFP fls2* with 1.25uM flg22 – overview

Movie 6: Treatment of *UBQ10::R-GECO1* in WT background with 1.25uM flg22 – zoom in elongation zone

Movie 7: Treatment of *UBQ10::R-GECO1 WER::FLS2-GFP fls2* with 1.25uM flg22 – zoom in elongation zone

Movie 8: Treatment of *UBQ10::R-GECO1 CASP1::FLS2-GFP fls2* with 1.25uM flg22 – zoom in differentiated zone

Movie 9: Treatment of *UBQ10::R-GECO1 SHR::FLS2-GFP fls2* with 1.25uM flg22 – zoom in differentiated zone

Movie 10: Treatment of *UBQ10::R-GECO1 UBQ10::FLS2-GFP fls2* with 1.25uM flg22 – zoom in differentiated zone

## Notes

### Competing Interest Statement

The authors have declared no competing interest.

## References

Adams-Phillips, L., Briggs, A.G., and Bent, A.F. (2010). Disruption of poly(ADP-ribosyl)ation mechanisms alters responses of *Arabidopsis* to biotic stress. Plant Physiol. 152, 267–280.

Alassimone, J., Fujita, S., Doblas, V.G., Dop, M. van, Barberon, M., Kalmbach, L., Vermeer, J.E.M., Rojas-Murcia, N., Santuari, L., Hardtke, C.S., et al. (2016). Polarly localized kinase SGN1 is required for Casparian strip integrity and positioning. Nat. Plants 2, 1–10.

Andersen, T.G., Barberon, M., and Geldner, N. (2015). Suberization — the second life of an endodermal cell. Curr. Opin. Plant Biol. 28, 9–15.

Andersen, T.G., Naseer, S., Ursache, R., Wybouw, B., Smet, W., De Rybel, B., Vermeer, J.E.M., and Geldner, N. (2018). Diffusible repression of cytokinin signalling produces endodermal symmetry and passage cells. Nature 555, 529–533.

Arora, S.K., Ritchings, B.W., Almira, E.C., Lory, S., and Ramphal, R. (1997). A transcriptional activator, FleQ, regulates mucin adhesion and flagellar gene expression in *Pseudomonas aeruginosa* in a cascade manner. J. Bacteriol. 179, 5574–5581.

Bai, Y., Müller, D.B., Srinivas, G., Garrido-Oter, R., Potthoff, E., Rott, M., Dombrowski, N., Münch, P.C., Spaepen, S., Remus-Emsermann, M., et al. (2015). Functional overlap of the *Arabidopsis* leaf and root microbiota. Nature 528, 364–369.

Beck, M., Wyrsch, I., Strutt, J., Wimalasekera, R., Webb, A., Boller, T., and Robatzek, S. (2014). Expression patterns of FLAGELLIN SENSING 2 map to bacterial entry sites in plant shoots and roots. J. Exp. Bot. 65, 6487–6498.

Benfey, P.N., Linstead, P.J., Roberts, K., Schiefelbein, J.W., Hauser, M.T., and Aeschbacher, R.A. (1993). Root development in *Arabidopsis*: four mutants with dramatically altered root morphogenesis. Development 119, 57–70.

Berendsen, R.L., Pieterse, C.M.J., and Bakker, P.A.H.M. (2012). The rhizosphere microbiome and plant health. Trends Plant Sci. 17, 478–486.

Bernards, M.A. (2002). Demystifying suberin. Can. J. Bot. 80, 227–240.

Bolte, S., and Cordelières, F.P. (2006). A guided tour into subcellular colocalization analysis in light microscopy. J. Microsc. 224, 213–232.

Bulgarelli, D., Schlaeppi, K., Spaepen, S., van Themaat, E.V.L., and Schulze-Lefert, P. (2013). Structure and Functions of the Bacterial Microbiota of Plants. Annu. Rev. Plant Biol. 64, 807–838.

Buscaill, P., Chandrasekar, B., Sanguankiattichai, N., Kourelis, J., Kaschani, F., Thomas, E.L., Morimoto, K., Kaiser, M., Preston, G.M., and Ichinose, Y. (2019). Glycosidase and glycan polymorphism control hydrolytic release of immunogenic flagellin peptides. PLANT Sci. 12.

Chezem, W.R., Memon, A., Li, F.-S., Weng, J.-K., and Clay, N.K. (2017). SG2-type R2R3-MYB transcription factor MYB15 controls defense-induced lignification and basal immunity in *Arabidopsis*. Plant Cell Online tpc.00954.2016.

Chinchilla, D., Zipfel, C., Robatzek, S., Kemmerling, B., Nürnberger, T., Jones, J.D.G., Felix, G., and Boller, T. (2007). A flagellin-induced complex of the receptor FLS2 and BAK1 initiates plant defence. Nature 448, 497–500.

Clough, S.J., and Bent, A.F. (1998). Floral dip: a simplified method for *Agrobacterium*-mediated transformation of *Arabidopsis thaliana*. Plant J. 16, 735–743.

Couto, D., and Zipfel, C. (2016). Regulation of pattern recognition receptor signalling in plants. Nat. Rev. Immunol. 16, 537.

De Coninck, B., Timmermans, P., Vos, C., Cammue, B.P.A., and Kazan, K. (2015). What lies beneath: belowground defense strategies in plants. Trends Plant Sci. 20, 91–101.

Doblas, V.G., Smakowska-Luzan, E., Fujita, S., Alassimone, J., Barberon, M., Madalinski, M., Belkhadir, Y., and Geldner, N. (2017). Root diffusion barrier control by a vasculature-derived peptide binding to the SGN3 receptor. Science 355, 280–284.

Dubiella, U., Seybold, H., Durian, G., Komander, E., Lassig, R., Witte, C.-P., Schulze, W.X., and Romeis, T. (2013). Calcium-dependent protein kinase/NADPH oxidase activation circuit is required for rapid defense signal propagation. Proc. Natl. Acad. Sci. 110, 8744–8749.

Faulkner, C., and Robatzek, S. (2012). Plants and pathogens: putting infection strategies and defence mechanisms on the map. Curr. Opin. Plant Biol. 15, 699–707.

Felix, G., Duran, J.D., Volko, S., and Boller, T. (1999). Plants have a sensitive perception system for the most conserved domain of bacterial flagellin. Plant J. 18, 265–276.

Fendrych, M., Van Hautegem, T., Van Durme, M., Olvera-Carrillo, Y., Huysmans, M., Karimi, M., Lippens, S., Guérin, C.J., Krebs, M., Schumacher, K., et al. (2014). Programmed cell death controlled by ANAC033/SOMBRERO determines root cap organ size in *Arabidopsis*. Curr. Biol. 24, 931–940.

Frangi, A.F., Niessen, W.J., Vincken, K.L., and Viergever, M.A. (1998). Multiscale vessel enhancement filtering. In Medical Image Computing and Computer-Assisted Intervention — MICCAI’98, W.M. Wells, A. Colchester, and S. Delp, eds. (Berlin, Heidelberg: Springer), pp. 130–137.

Fujita, S., De Bellis, D., Edel, K.H., Köster, P., Andersen, T.G., Schmid-Siegert, E., Dénervaud Tendon, V., Pfister, A., Marhavý, P., Ursache, R., et al. (2020). SCHENGEN receptor module drives localized ROS production and lignification in plant roots. EMBO J. n/a, e103894.

Garrido-Oter, R., Nakano, R.T., Dombrowski, N., Ma, K.-W., McHardy, A.C., and Schulze-Lefert, P. (2018). Modular traits of the rhizobiales root microbiota and their evolutionary relationship with symbiotic rhizobia. Cell Host Microbe 24, 155–167.e5.

Gilroy, S., Suzuki, N., Miller, G., Choi, W.-G., Toyota, M., Devireddy, A.R., and Mittler, R. (2014). A tidal wave of signals: calcium and ROS at the forefront of rapid systemic signaling. Trends Plant Sci. 19, 623–630.

Gilroy, S., Białasek, M., Suzuki, N., Górecka, M., Devireddy, A., Karpinski, S., and Mittler, R. (2016). ROS, calcium and electric signals: key mediators of rapid systemic signaling in plants. Plant Physiol. pp.00434.2016.

Gómez-Gómez, L., and Boller, T. (2000). FLS2: An LRR receptor–like kinase involved in the perception of the bacterial elicitor flagellin in *Arabidopsis*. Mol. Cell 5, 1003–1011.

Gómez-Gómez, L., Felix, G., and Boller, T. (1999). A single locus determines sensitivity to bacterial flagellin in *Arabidopsis thaliana*. Plant J. 18, 277–284.

Gul-Mohammed, J., Arganda-Carreras, I., Andrey, P., Galy, V., and Boudier, T. (2014). A generic classification-based method for segmentation of nuclei in 3D images of early embryos. BMC Bioinformatics 15, 9.

Hara, M., Umetsu, N., Miyamoto, C., and Tamari, K. (1973). Inhibition of the biosynthesis of plant cell wall materials, especially cellulose biosynthesis, by coumarin. Plant Cell Physiol. 14, 11–28.

Helariutta, Y., Fukaki, H., Wysocka-Diller, J., Nakajima, K., Jung, J., Sena, G., Hauser, M.-T., and Benfey, P.N. (2000). The SHORT-ROOT gene controls radial patterning of the *Arabidopsis* root through radial signaling. Cell 101, 555–567.

Hijwegen, T. (1963). Lignification, a possible mechanism of active resistance against pathogens. Neth. J. Plant Pathol. 69, 314–317.

Jeworutzki, E., Roelfsema, M.R.G., Anschütz, U., Krol, E., Elzenga, J.T.M., Felix, G., Boller, T., Hedrich, R., and Becker, D. (2010). Early signaling through the *Arabidopsis* pattern recognition receptors FLS2 and EFR involves Ca2+-associated opening of plasma membrane anion channels. Plant J. 62, 367–378.

Kamula, S.A., Peterson, C.A., and Mayfield, C.I. (1994). Impact of the exodermis on infection of roots by *Fusarium culmorum*. Plant Soil 167, 121–126.

Keinath, N.F., Waadt, R., Brugman, R., Schroeder, J.I., Grossmann, G., Schumacher, K., and Krebs, M. (2015). Live cell imaging with R-GECO1 sheds light on flg22- and chitin-induced transient [Ca2+]cyt patterns in *Arabidopsis*. Mol. Plant 8, 1188–1200.

Kumpf, R.P., and Nowack, M.K. (2015). The root cap: a short story of life and death. J. Exp. Bot. 66, 5651–5662.

Kupferschmied, P. (2015). Molecular basis and regulation of insect pathogenicity in plant-beneficial pseudomonads. Université de Lausanne, Faculté de biologie et médecine.

Kupferschmied, P., Péchy-Tarr, M., Imperiali, N., Maurhofer, M., and Keel, C. (2014). Domain shuffling in a sensor protein contributed to the evolution of insect pathogenicity in plant-beneficial *Pseudomonas protegens*. PLOS Pathog. 10, e1003964.

Kurihara, D., Mizuta, Y., Sato, Y., and Higashiyama, T. (2015). ClearSee: a rapid optical clearing reagent for whole-plant fluorescence imaging. Development 142, 4168–4179.

Lange, B.M., Lapierre, C., and Jr, H.S. (1995). Elicitor-induced spruce stress lignin (structural similarity to early developmental lignins). Plant Physiol. 108, 1277–1287.

Lee, M.M., and Schiefelbein, J. (1999). WEREWOLF, a MYB-related protein in *Arabidopsis*, is a position-dependent regulator of epidermal cell patterning. Cell 99, 473–483.

Lee, M.-H., Jeon, H.S., Kim, S.H., Chung, J.H., Roppolo, D., Lee, H.-J., Cho, H.J., Tobimatsu, Y., Ralph, J., and Park, O.K. (2019). Lignin-based barrier restricts pathogens to the infection site and confers resistance in plants. EMBO J. 38, e101948.

Li, B., Meng, X., Shan, L., and He, P. (2016). Transcriptional Regulation of Pattern-Triggered Immunity in Plants. Cell Host Microbe 19, 641–650.

Mandal, S., and Mitra, A. (2007). Reinforcement of cell wall in roots of *Lycopersicon esculentum* through induction of phenolic compounds and lignin by elicitors. Physiol. Mol. Plant Pathol. 71, 201– 209.

Marhavý, P., Kurenda, A., Siddique, S., Dénervaud Tendon, V., Zhou, F., Holbein, J., Hasan, M.S., Grundler, F.M., Farmer, E.E., and Geldner, N. (2019). Single-cell damage elicits regional, nematode-restricting ethylene responses in roots. EMBO J. 38, e100972.

Marquès-Bueno, M.M., Morao, A.K., Cayrel, A., Platre, M.P., Barberon, M., Caillieux, E., Colot, V., Jaillais, Y., Roudier, F., and Vert, G. (2015). A versatile Multisite Gateway-compatible promoter and transgenic line collection for cell type-specific functional genomics in *Arabidopsis*. Plant J. n/a-n/a.

Martínez-García, E., and Lorenzo, V. de (2011). Engineering multiple genomic deletions in Gram-negative bacteria: analysis of the multi-resistant antibiotic profile of *Pseudomonas putida* KT2440. Environ. Microbiol. 13, 2702–2716.

Messner, B., and Boll, M. (1993). Elicitor-mediated induction of enzymes of lignin biosynthesis and formation of lignin-like material in a cell suspension culture of spruce (*Picea abies*). Plant Cell Tissue Organ Cult. 34, 261–269.

Millet, Y.A., Danna, C.H., Clay, N.K., Songnuan, W., Simon, M.D., Werck-Reichhart, D., and Ausubel, F.M. (2010). Innate Immune Responses Activated in *Arabidopsis* Roots by Microbe-Associated Molecular Patterns. Plant Cell 22, 973–990.

Nicholson, R.L., and Hammerschmidt, R. (1992). Phenolic Compounds and Their Role in Disease Resistance. Annu. Rev. Phytopathol. 30, 369–389.

Ollion, J., Cochennec, J., Loll, F., Escudé, C., and Boudier, T. (2013). TANGO: a generic tool for high-throughput 3D image analysis for studying nuclear organization. Bioinformatics 29, 1840–1841.

Pel, M.J.C., and Pieterse, C.M.J. (2013). Microbial recognition and evasion of host immunity. J. Exp. Bot. 64, 1237–1248.

Pfister, A., Barberon, M., Alassimone, J., Kalmbach, L., Lee, Y., Vermeer, J.E., Yamazaki, M., Li, G., Maurel, C., Takano, J., et al. (2014). A receptor-like kinase mutant with absent endodermal diffusion barrier displays selective nutrient homeostasis defects. ELife 3, e03115.

Poncini, L., Wyrsch, I., Tendon, V.D., Vorley, T., Boller, T., Geldner, N., Métraux, J.-P., and Lehmann, S. (2017). In roots of *Arabidopsis thaliana*, the damage-associated molecular pattern AtPep1 is a stronger elicitor of immune signalling than flg22 or the chitin heptamer. PLOS ONE 12, e0185808.

Ranathunge, K., Thomas, R.H., Fang, X., Peterson, C.A., Gijzen, M., and Bernards, M.A. (2008). Soybean Root Suberin and Partial Resistance to Root Rot Caused by *Phytophthora sojae*. Phytopathology 98, 1179–1189.

Rich-Griffin, C., Eichmann, R., Reitz, M.U., Hermann, S., Woolley-Allen, K., Brown, P.E., Wiwatdirekkul, K., Esteban, E., Pasha, A., Kogel, K.-H., et al. (2020). Regulation of Cell Type-Specific Immunity Networks in *Arabidopsis* Roots. Plant Cell.

Robatzek, S., Chinchilla, D., and Boller, T. (2006). Ligand-induced endocytosis of the pattern recognition receptor FLS2 in *Arabidopsis*. Genes Dev. 20, 537–542.

Robertsen, B. (1986). Elicitors of the production of lignin-like compounds in cucumber hypocotyls. Physiol. Mol. Plant Pathol. 28, 137–148.

Schneider, C.A., Rasband, W.S., and Eliceiri, K.W. (2012). NIH Image to ImageJ: 25 years of image analysis. Nat. Methods 9, 671–675.

Seybold, H., Trempel, F., Ranf, S., Scheel, D., Romeis, T., and Lee, J. (2014). Ca2+ signalling in plant immune response: from pattern recognition receptors to Ca^2+^ decoding mechanisms - Seybold – 2014 - - Wiley Online Library. New Phytol.

Siligato, R., Wang, X., Yadav, S.R., Lehesranta, S., Ma, G., Ursache, R., Sevilem, I., Zhang, J., Gorte, M., Prasad, K., et al. (2016). MultiSite Gateway-Compatible Cell Type-Specific Gene-Inducible System for Plants. Plant Physiol. 170, 627–641.

Smit, F., and Dubery, I.A. (1997). Cell wall reinforcement in cotton hypocotyls in response to a *Verticillium dahliae* elicitor. Phytochemistry 44, 811–815.

Stanley, C.E., Shrivastava, J., Brugman, R., Heinzelmann, E., Swaay, D. van, and Grossmann, G. (2018). Dual-flow-RootChip reveals local adaptations of roots towards environmental asymmetry at the physiological and genetic levels. New Phytol. 217, 1357–1369.

Stringlis, I.A., de Jonge, R., and Pieterse, C.M.J. (2019). The Age of Coumarins in Plant–Microbe Interactions. Plant Cell Physiol.

Stutz, E.W., Défago, G., and Kern, H. (1986). Naturally occurring fluorescent pseudomonads involved in suppression of black root rot of tobacco. Phytopathology 76, 181–185.

Tang, D., Wang, G., and Zhou, J.-M. (2017). Receptor Kinases in Plant-Pathogen Interactions: More Than Pattern Recognition. Plant Cell 29, 618–637.

Thomas, R., Fang, X., Ranathunge, K., Anderson, T.R., Peterson, C.A., and Bernards, M.A. (2007). Soybean Root Suberin: Anatomical Distribution, Chemical Composition, and Relationship to Partial Resistance to *Phytophthora sojae*. Plant Physiol. 144, 299–311.

Thor, K., and Peiter, E. (2014). Cytosolic calcium signals elicited by the pathogen-associated molecular pattern flg22 in stomatal guard cells are of an oscillatory nature. New Phytol. 204, 873–881.

Tognolli, M., Penel, C., Greppin, H., and Simon, P. (2002). Analysis and expression of the class III peroxidase large gene family in *Arabidopsis thaliana*. Gene 288, 129–138.

Ursache, R., Andersen, T.G., Marhavý, P., and Geldner, N. (2018). A protocol for combining fluorescent proteins with histological stains for diverse cell wall components. Plant J. 93, 399–412.

Vance, C.P., Kirk, T.K., and Sherwood, R.T. (1980). Lignification as a Mechanism of Disease Resistance. Annu. Rev. Phytopathol. 18, 259–288.

Vermeer, J.E.M., Wangenheim, D. von, Barberon, M., Lee, Y., Stelzer, E.H.K., Maizel, A., and Geldner, N. (2014). A Spatial Accommodation by Neighboring Cells Is Required for Organ Initiation in *Arabidopsis*. Science 343, 178–183.

Wyrsch, I., Domínguez-Ferreras, A., Geldner, N., and Boller, T. (2015). Tissue-specific FLAGELLIN-SENSING 2 (FLS2) expression in roots restores immune responses in *Arabidopsis* fls2 mutants. New Phytol. 206, 774–784.

Yamaguchi, S., Fujita, H., Sugata, K., Taira, T., and Iino, T. (1984). Genetic Analysis of H2, the Structural Gene for Phase-2 Flagellin in *Salmonella*. Microbiology, 130, 255–265.

Yu, K., Pieterse, C.M.J., Bakker, P.A.H.M., and Berendsen, R.L. (2019a). Beneficial microbes going underground of root immunity. Plant Cell Environ. 42, 2860–2870.

Yu, K., Liu, Y., Tichelaar, R., Savant, N., Lagendijk, E., Kuijk, S.J.L. van, Stringlis, I.A., Dijken, A.J.H. van, Pieterse, C.M.J., Bakker, P.A.H.M., et al. (2019b). Rhizosphere-Associated *Pseudomonas* Suppress Local Root Immune Responses by Gluconic Acid-Mediated Lowering of Environmental pH. Curr. Biol. 0.

Zhou, F., Emonet, A., Dénervaud Tendon, V., Marhavy, P., Wu, D., Lahaye, T., and Geldner, N. (2020). Co-incidence of Damage and Microbial Patterns Controls Localized Immune Responses in Roots. Cell 180, 440–453.e18.

Zipfel, C. (2008). Pattern-recognition receptors in plant innate immunity. Curr. Opin. Immunol. 20, 10– 16.

Zipfel, C., Robatzek, S., Navarro, L., Oakeley, E.J., Jones, J.D.G., Felix, G., and Boller, T. (2004). Bacterial disease resistance in *Arabidopsis* through flagellin perception. Nature 428, 764–767.

